# Multi-Dimensional Machine Learning Approaches for Fruit Shape Recognition and Phenotyping in Strawberry

**DOI:** 10.1101/736397

**Authors:** Mitchell J. Feldmann, Michael A. Hardigan, Randi A. Famula, Cindy M. López, Amy Tabb, Glenn S. Cole, Steven J. Knapp

## Abstract

**Background:** Shape is a critical element of the visual appeal of strawberry fruit and determined by both genetic and non-genetic factors. Current fruit phenotyping approaches for external characteristics in strawberry rely on the human eye to make categorical assessments. However, fruit shape is multi-dimensional, continuously variable, and not adequately described by a single quantitative variable. Morphometric approaches enable the study of complex forms but are often abstract and difficult to interpret. In this study, we developed a mathematical approach for transforming fruit shape classifications from digital images onto an ordinal scale called the principal progression of k clusters (PPKC). We use these human-recognizable shape categories to select features extracted from multiple morphometric analyses that are best fit for genome-wide and forward genetic analyses.

**Results:** We transformed images of strawberry fruit into human-recognizable categories using unsupervised machine learning, discovered four principal shape categories, and inferred progression using PPKC. We extracted 67 quantitative features from digital images of strawberries using a suite of morphometric analyses and multi-variate approaches. These analyses defined informative feature sets that effectively captured quantitative differences between shape classes. Classification accuracy ranged from 68.9 – 99.3% for the newly created, genetically correlated phenotypic variables describing a shape.

**Conclusions:** Our results demonstrated that strawberry fruit shapes could be robustly quantified, accurately classified, and empirically ordered using image analyses, machine learning, and PPKC. We generated a dictionary of quantitative traits for studying and predicting shape classes and identifying genetic factors underlying phenotypic variability for fruit shape in strawberry. The methods and approaches we applied in strawberry should apply to other fruits, vegetables, and specialty crops.

## Background

Fruit breeders actively selected several morphological and quality phenotypes during the domestication of garden strawberry (*Fragaria* × *ananassa*), an allooctoploid (2n = 8x = 56) of hybrid origin [1, 2, 3]. *F*. × *ananassa* was created in the early 1700s by interspecific hybridization between ecotypes of wild octoploid species (*F. virginiana* and *F. chiloensis*), multiple subsequent introgressions of genetic diversity from *F. virginiana* and *F. chiloensis* subspecies in subsequent generations, and artificial selection for horticulturally important traits among interspecific hybrid descendants. Domestication and breeding have altered the fruit morphology, development, and metabolome of garden strawberry, distancing modern cultivars from their wild progenitors [4, 5, 6, 7, 8, 9]. Approximately 300 years of breeding in the admixed hybrid population has led to the emergence of high yielding cultivars with large, firm, visually appealing, long shelf-life fruit that can withstand the rigors of harvest, handling, storage, and long-distance shipping [10]. Fruit shape is an essential trait of agricultural products, particularly those of specialty crops, due to perceived and realized relationships with the quality and value of the products. Image-based plant phenotyping has the potential to increase scope, throughput, and accuracy in forward-genetic studies by reducing the effects of user bias, enabling the analysis of larger sample sizes, and partitioning of genetic variance from other environments (E), management (M), and other non-genetic sources of variation [11, 12, 13].

Current fruit phenotyping approaches for external characteristics in strawberry rely on the human eye to make categorical assessments [14, 15, 16]. Descriptive categories for planar shapes (e.g., rhombic and reniform) have long played a role in plant systematics [17]. Categories may be either nominal [11, 18, 19], existing in name only, or ordinal, referring to a position in an ordered series or on a gradient [15, 16, 19]. Classification is often labor-intensive and prone to human bias, which can increase with task complexity and time requirement [20, 21]. Alternative scoring approaches have relied on morphometrics and machine learning to automate classification; for example, sorting fruit into shape categories in both tomato [11] and strawberry [18]. Unsupervised machine learning methods (e.g., *k*-means, hierarchical, and Bayesian clustering), unlike supervised methods, are useful for pattern detection and clustering, while supervised machine learning methods (e.g., regression, discriminant analysis, and support vector regression) are useful for prediction and classification [22, 23]. Unsupervised clustering enables the calculation of several measures of model performance and overfitting to balance compression and accuracy. However, the categories derived from these techniques are without order, resulting in the need for a suitable transformation to an ordinal scale, which is more appropriate for quantitative genetic analyses [24, 25, 26, 27, 28]. In this context, ordinal categories give the interpretation of relationship with, or distance from, other shape categories in the series. To enable this interpretation, we developed a method for discovering the progression through fruit shape categories derived from unsupervised machine learning methods. The principal progression of *k* clusters (PPKC), allowed us to non-arbitrarily determine the appropriate shape gradient for statistical analyses using empirical data. Here, we describe approaches for translating digital images of strawberries into computationally defined phenotypic variables for identifying and classifying fruit shapes.

Fruit shape and anatomy are complex, multi-dimensional, and abstract phenotypes that are often not readily or intuitively described by planar descriptors or individual qualitative or quantitative phenotypes. Beyond precise qualitative definitions used in systematics (e.g., rhombic, falcate, and reniform) [17, 18], references to fruit shape encompass a wide variety of mathematical parameters and geometric indices that establish quantitative measurements of plant organs [29, 30, 31]. Much like human faces, fruit shape and anatomy are products of the underlying genetic and non-genetic determinants of phenotypic variability in a population [32, 33]. The genetic determinants of fruit shape are unknown in strawberry, in part because researchers have not yet translated fruit shape attributes into quantitative phenotypic variables, which are essential for identifying the underlying genes or quantitative trait loci through genome-wide association studies (GWAS) and other forward-genetic approaches [34, 35, 36, 37]. Quantitative phenotypic measurements have allowed researchers to uncover some of the genetic basis of fruit shape in tomato [38, 39], pepper [40, 41], pear [42], melon [33], potato [43], and strawberry [9, 44]. These quantitative features often rely on linear metrics of distance (e.g., height, width, and perimeter) and are generally modified into compound descriptors that remove the effects of size (e.g., aspect ratio or roundness) [42, 44, 45]. However, compound linear descriptors often have limited resolution compared to more comprehensive, multi-variate descriptors [31]. Elliptical Fourier Analysis (EFA) quantifies fruit shape from a closed outline by converting a closed-contour into a weighted sum of wave functions with different frequencies [12, 46, 47, 48, 49, 50]. Generalized Procrustes Analysis (GPA) quantifies the distance between sets of biologically homologous, or mathematically similar, (pseudo-)landmarks on the surface of an object [49, 51, 52, 53, 54, 55, 56]. Fruit shape can also be described using linear combinations of pixel intensities from digital images extrapolating from analyses generally used to quantify color patterns and facial recognition [13, 57, 58, 59, 60, 61, 62]. Here, we generated a dictionary of 67 quantitative features, including linear-, outline-, landmark-, and pixel-based descriptors to investigate the quality of different features in preparation for forward-genetic analyses.

The ultimate goal of our study was to develop heritable phenotypic variables for describing fruit shape, which could then be used to identify the genetic factors underlying phenotypic differences in fruit shape. We describe and demonstrate the application of PPKC, which transforms categories discovered from unsupervised machine learning methods to a more convenient and analytically tractable ordinal scale [24, 26, 27]. We explore the relationship between machine-acquired categories and 67 quantitative features extracted from digital images. We apply random forest regression to select critical sets of quantitative features for classification and use supervised machine learning methods, including support vector regression and linear discriminant analysis, to determine the accuracy of shape classification. We discovered that there are only a few categories of interest in a highly domesticated breeding population and that a small number of features are needed to classify shape into the discovered categories accurately. We also find that ordinal shape categories are highly heritable and that the features needed for accurate classification are also heritable.

## Data Description

The data released with this manuscript contains digital images of 6, 874 strawberry fruit from 572 hybrids originating from the University of California, Davis Strawberry Breeding Program. The data for this manuscript, including pre-processed images (Fig. 1A), processed images (Fig. 1B), and extracted features (see Methods), are available on Zenodo [63]. The pre-processed images typically contained multiple berries per image along with a data matrix bar code indicating the genotype ID and other elements of the experiment design. The processed images are 1000 × 1000px-scaled binary images of individual fruit. The extracted features data set is provided as a CSV file. The code to replicate the analyses in this manuscript is provided in a GitHub repository [64]. We hope that the release of this data assists others in developing novel morphometric approaches to better understand the genetic, developmental, and environmental control of fruit shape in strawberry, and more broadly in other fruits, vegetables, and specialty crops.

**Figure 1.**
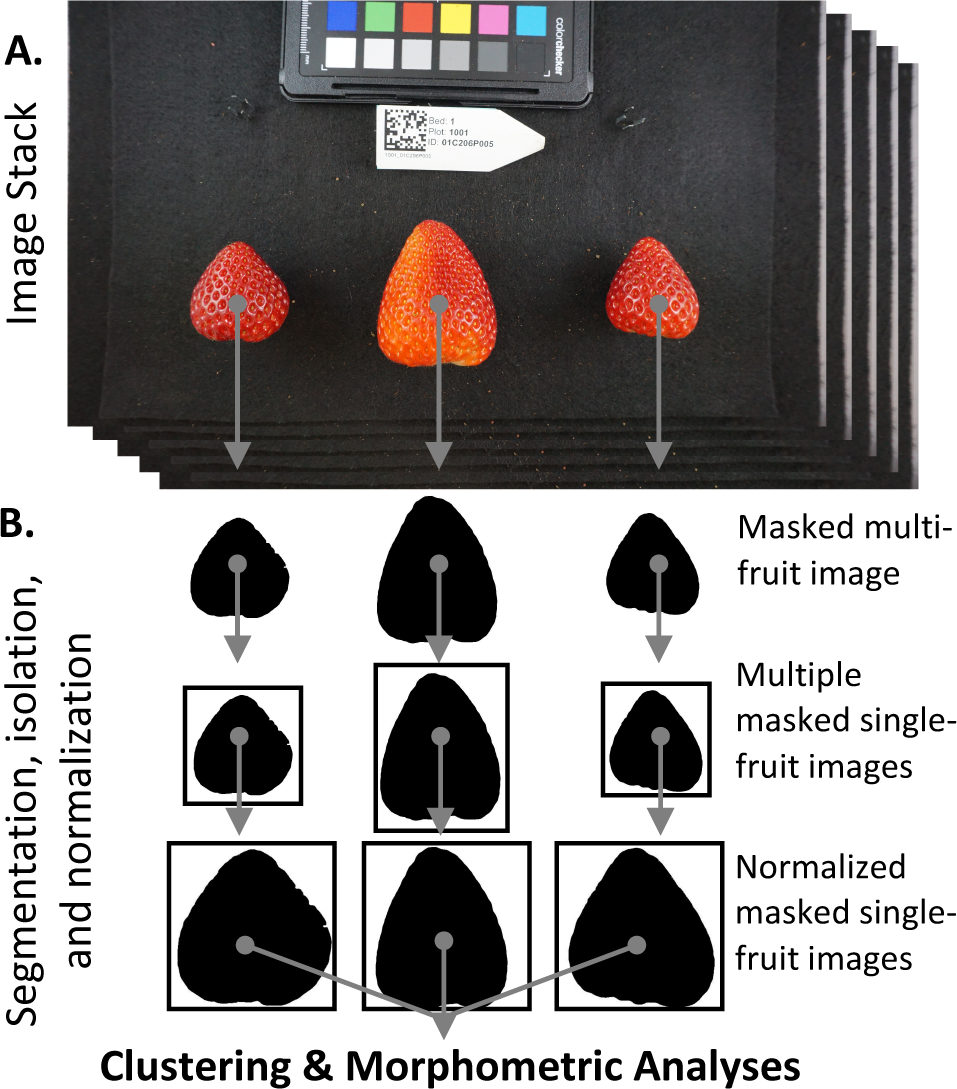
An example of the processing pipeline. **(A)** A user collects a stack of images containing multiple strawberries and a unique QR code. **(B)** All images are then segmented using the SIOX algorithm implemented in ImageJ. Each object is then cut from its original image based on the coordinates of its bounding rectangle in R 3.5.3. White pixels are then added to the edges of each frame until all images are 1000 × 1000 pixels. Regions of interest are then scaled such that the major axis of each object becomes 1000px in ImageJ. Output images are scale invariant and maintain the original aspect ratio.

## Analyses

### Modified k-means clustering

*k*-means clustering can rapidly detect patterns in large, multi-dimensional data sets used for clustering, decision making, and dimension reduction [22, 65, 66]. It is an iterative algorithm that partitions a data set into a pre-defined number of non-overlapping clusters, *k*, by minimizing the sum of squared distances from each data point to the cluster centroid. A centroid corresponds to the mean of all points assigned to the cluster. Here, we used *k*-means to cluster flattened binary images (Fig. 1; see Methods). Individual fruits were segmented from the image background as a binary mask, normalized by the major axis, resized to 100 × 100px, and flattened into a vector (Fig. 1; see Methods). We represented each image as a 10, 000 element vector containing binary pixel values. We were able to rapidly and reliably assign images to classes using *k*-means clustering. In this experiment, we allowed *k*, the number of permitted categories, to range from 2 to 10. We anticipated that a human-based classification system would not have the speed or reliability needed for this task, particularly for larger values of *k*. We visualized the centroids for each class (Fig. S1A). Several groups were found to be mirror images of one another when *k* = 8 (Fig. S1B, black squares). The level plots in Figure S1 depict shape outlines reflecting the 20^th^, 40^th^, 60^th^, and 80^th^ quantiles. To check mirror symmetry, we first rotated one of the suspect classes about the vertical axis (the proposed axis of symmetry) (Fig. S1C; dark gray square) and then overlaid the rotated centroid onto the alternate, unrotated centroid (Fig. S1D). This type of symmetry in the clusters is assumed to have arisen from orientation artifacts during imaging (Fig. 1). It seemed likely that if we observed an object from two opposite sides, they would appear as mirror images of one another and would not reflect an actual difference in category. In this example, 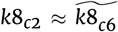, where 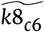 is the rotated *k*8_*c*6_ centroid. *k*8 refers to the results of clustering with *k* = 8 groups, whereas *c*2 and *c*6 refer to the cluster assignment within a value of *k*. The Euclidean distance between the centroid of *k*8_*c*2_ and 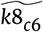was 6.93, and the Euclidean distance between the centroid of *k*8_*c*2_ and *k*8_*c*6_ was 15.73. We re-ran *k*-means clustering but replaced all images in *k*8_*c*6_ with those in 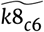. The unrotated centroid *k*8_*c*6_ was 2.26× more dissimilar to *k*8_*c*2_ than the rotated centroid 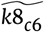. As intended, the rotation of *k*8_*c*6_ collapsed the two clusters with reflective symmetry into one and exposed a new cluster (Fig. S1E; light gray square).

### Principal progression of k clusters

*k*-means clustering does not assign a progression or gradient to discovered classes. However, score and ordinal traits are typically more useful in genetic studies than variables on nominal scales [24]. We developed a new method to transform the categories derived from *k*-means onto an ordinal scale, which we call the principal progression of *k* clusters, or PPKC (Fig. 2; Alg. 1). This method relies on *k*-means clustering to categorize images. The *k*-means analysis supports several metrics for evaluating model performance and overfit, including adjusted R^2^, AIC, and BIC, which allows users to determine the most appropriate value of *k* given the observed data. The gradient between clusters was estimated by performing principal components analysis on a covariance matrix reflecting the hierarchical relationship between a focal cluster and all previously discovered clusters.

**Figure 2.**
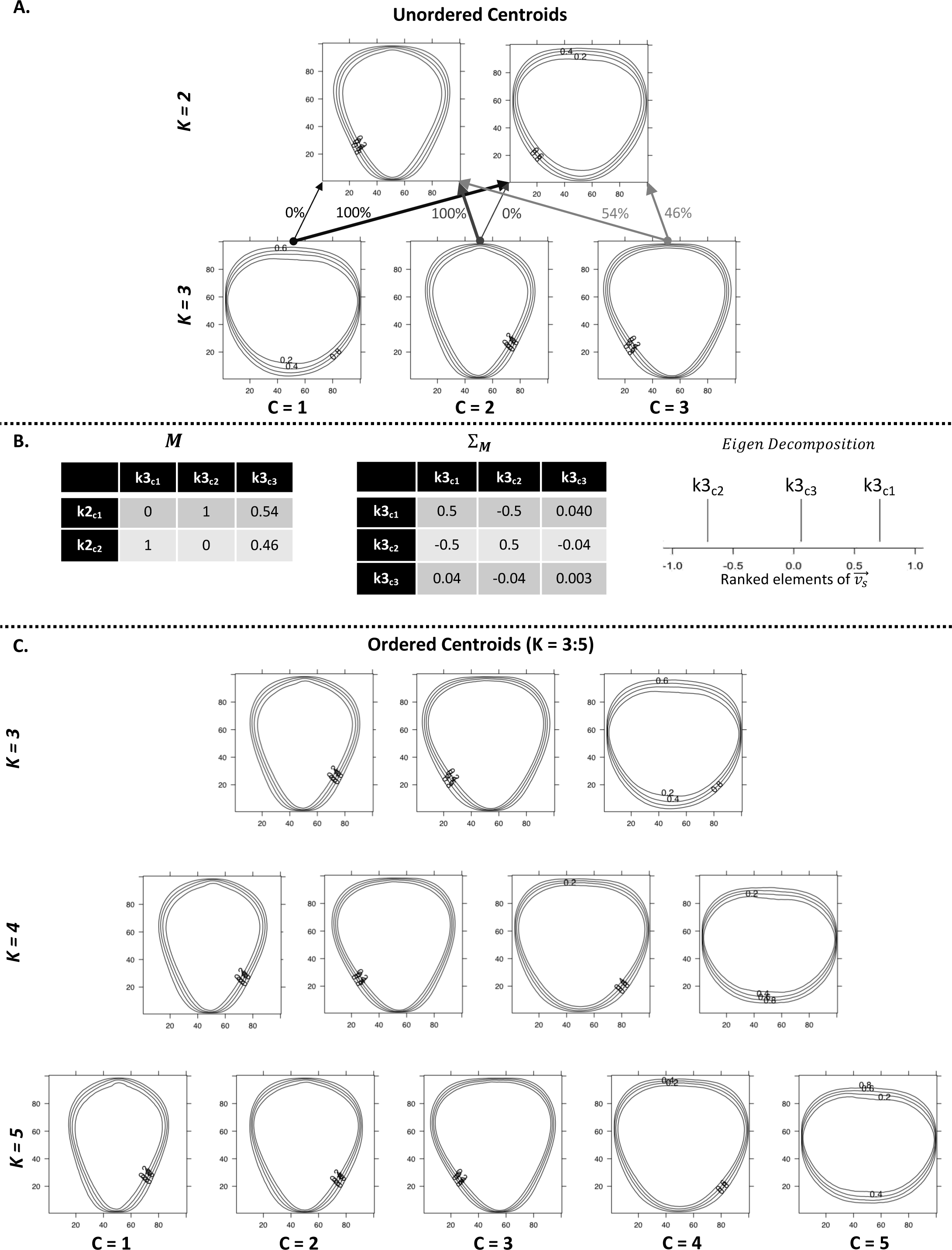
An example use of PPKC. **(A)** After *k*-means clustering is performed clusters are randomly assigned a numeric value (1,2,…,*k*). When *k* > 2, this value becomes nominal. PPKC relies on the fact that the order through clusters when *k* = 2 has identical interpretations in either direction. The lines representing each clusters centroid reflect the 20th, 40th, 60th, and 80th quantiles, moving out from the center of each images. **(B)**(1) A table representation of the resultant matrix from equation 1. Each cell represents the proportion of images in the column class and in the row class, normalized by the number of images in the column class. **(B)**(2) A table representation of Σ_**M**_. (B)(3) The ranked elements of 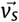 shown on a number line. (C) After using PPKC, the order of groups is explicitly identified. In this example, showing *k* = [3, 5], the order discovered seems to trend from tall and thin berries, through more triangular shapes, and ending with berries that are short and wide.

We first assigned each flattened binary image (Fig. 1) to a category using a modified *k*-means approach. We assigned a cluster to each image and allowed the number of clusters, *k*, to range from [2, 10]. The order was subsequently inferred using PPKC (Fig. 2; Alg. 1). When *k* = 2, the order of relatedness is arbitrary, and both *k*2_*c*1_ → *k*2_*c*2_ and *k*2_*c*2_ → *k*2_*c*1_ have the same meaning, where “→” indicates the progression of discovered categories. Any given order and its reverse are considered equivalent, and this applies to higher levels of *k* as well; for example, the hypothetical ranking 1, 4, 2, 3 is considered the equivalent of 3, 2, 4, 1. The phenotypic variance of two opposing ordinal scales (e.g., 1, 4, 2, 3) does not change. For each cluster of interest (e.g., *k*4_*c*1_, *k*4_*c*2_, *k*4_*c*3_, and *k*4_*c*4_), we calculated the proportion of each cluster that came from *k*3_*c*1_, *k*3_*c*2_, or *k*3_*c*3_ and *k*2_*c*1_ or *k*2_*c*2_ (i.e., all former classifications). These proportions enable the estimation of similarity between a cluster of interest (e.g., *k*4_*c*1_) and the clusters of all prior values of *k*. We then normalized the proportions by the total number of images in the cluster of interest (e.g., *k*4_*c*1_, *k*4_*c*2_, *k*4_*c*3_, and *k*4_*c*4_) (Eqn. 1).

For every level of *k* > 2, we constructed **M**, a rectangular matrix of size 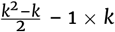 (Alg. 1 line 13). The sum of each column should equal *k* –2. The proportions were continuous values in the range [0, 1] that described the origin of a particular cluster of interest (e.g., *k*4_*c*1_) as it relates to the clusters of *k* = 3 and *k* = 2 or all clusters [2, *k* –1]. In this example, *k* = 4:

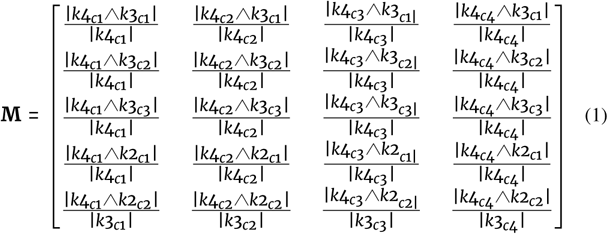

We then calculated the variance-covariance matrix of Eqn. (1) (Alg. 1; line 18). The variance-covariance matrix, Σ_**M**_, represents the relationship between each cluster of interest (e.g., *k*4_*c*1_, *k*4_*c*2_, *k*4_*c*3_,or *k*4_*c*4_).

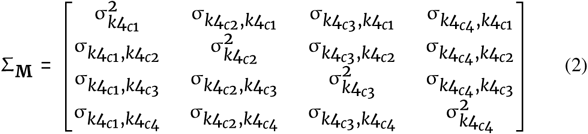

We then performed eigen decomposition on Eqn. (2) using the following equation (Alg. 1; line 19).

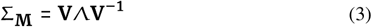

In Eqn. (3), Λ is a diagonal matrix with values corresponding to the *k* eigenvalues of Σ_**M**_ and **V** is a square matrix containing eigenvectors associated with the eigenvalues in Λ. We then extracted the eigenvector associated with the largest eigenvalue,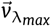. We ordered the elements of 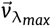 such that the resultant vector,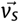, has the property *v*_*s*1_ *≤* … *≤ v*_*sk*_. We do not consider the distance between elements in 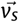, only their rank. The clusters are then indexed to match the rank of the associated elements in 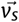. There are at most *k* eigenvalues associated with eigenvectors of length *k* due to Σ_**M**_ being *k* × *k*. Eigen decomposition is used to describe the major axis of variance in Σ_**M**_. In theory, this single-axis should be able to separate the classes more effectively than either the proportions or covariance measures.

After applying PPKC, the order of elements in 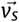 is taken to be the progression through the discovered categories. However, a single dimension may not capture the complexity of some relationships. In this study, we reached that limit when *k* = 8. The final three clusters in the progression (i.e., *k*8_*c*4_, *k*8_*c*7_, and *k*8_*c*8_) did not seem to follow the same pattern as in previous progressions (Fig. S2). The change in progression could be reflective of overfitting the number of groups in *k*-means clustering. The dramatic change of slope in the total within-group sums of squares, AIC, and Adjusted *R*^2^ evidenced overfitting (Fig. S3). The strongest evidence for four clusters is in the BIC, which is minimized when *k* = 4 (Fig. S3D). The elements of 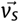 tend to converge on one another as *k* increases, which may be indicative of little biological information in the new clusters and overfitting (Fig. S4). We extrapolate that PPKC should continue to work beyond *k* = 8 if new clusters are biologically distinct and do not arise as an artifact of overfitting *k*. The order through categories was similar to those used in [14] and [16] and are characterized by a progression from ‘longer-than-wide’ (prolate) to ‘wider-than-long’ (oblate) (Fig. 2).

#### Algorithm 1 Principal Progression of K Clusters (PPKC) Algorithm

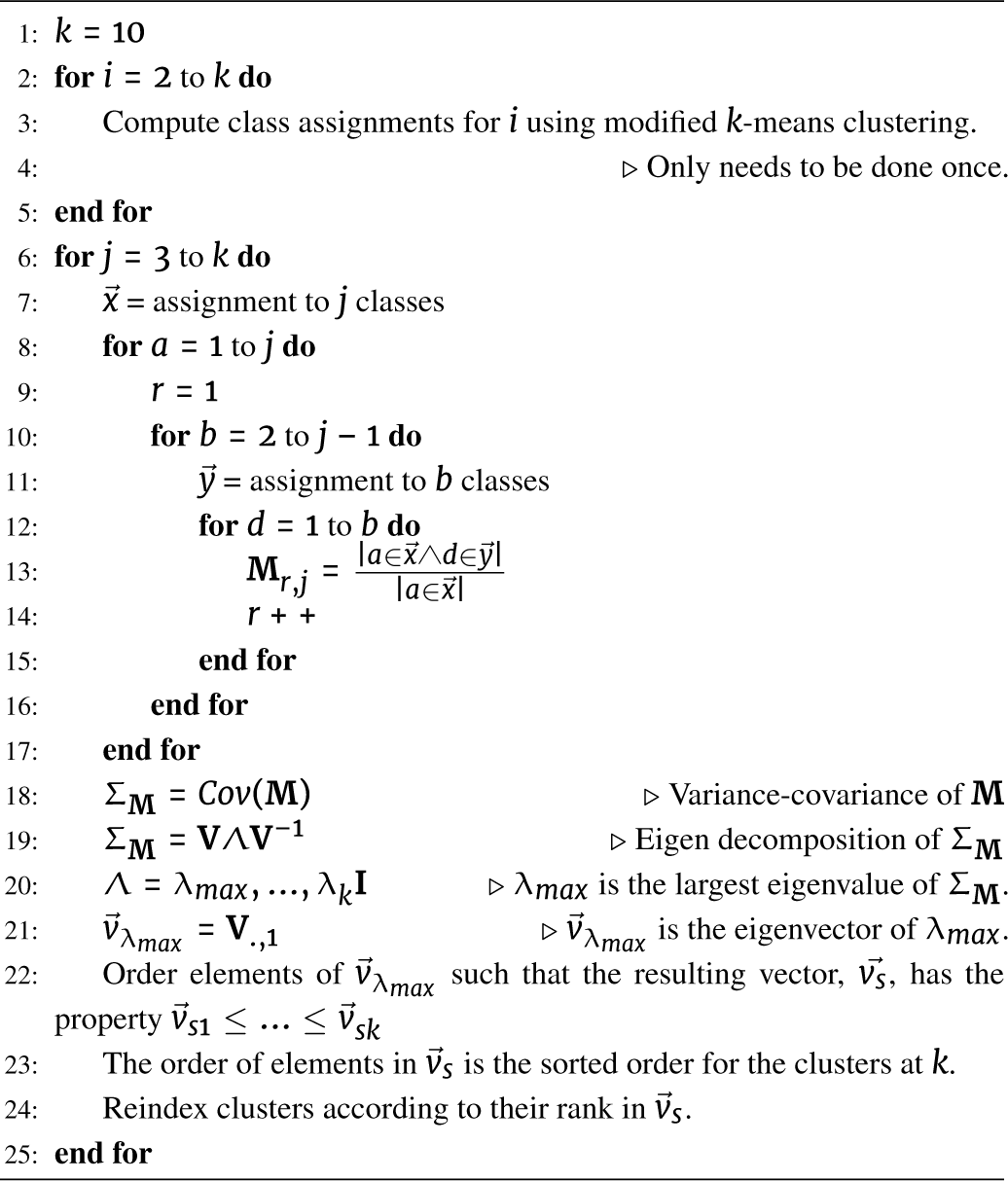

### Broad-sense heritability of ordered categories

For each value of *k*, broad-sense heritability (*H*^2^) on a clone-mean basis was assessed using a general linear mixed model with a cumulative logit link function (see Methods; Eqn. 8 and Eqn. 9) [67]. For this data set, *H*^2^ was generally high, ranging from *H*^2^ = 0.80 to 0.97, even as *k* → 10 (Table 2). These estimates of *H*^2^ are very similar to those reported in [16] (i.e., *H*^2^ = 0.84). When the *H*^2^ of a trait is in this range, it indicates that independent replications of the same individuals share a high degree of similarity and that most of the variation among individuals originated from genetic variation among individuals. Since the plant material used in this study are genetic clones, any variation in fruit shape among replicates originated from random, unobserved effects. For *k ≥* 8, the accuracy of *H*^2^ estimates is expected to be lower than for *k ≤* 7 as the gradient of the phenotype seems to be improperly specified. In this set of germplasm, we propose a set of four primary classes for categorizing fruit shape (Fig. 2 and S3). As *k* increases from 5 to 10, the visual similarity of some clusters is high (Fig. S2), thus indicating fewer relevant delineations (Fig. S4). As indicated, there is strong evidence in this data that there are four distinct clusters in this data (Fig S3).

### Feature selection using random forests

To discover which of 67 quantitative features (summarized in Figures 3 and 4) capture and reflect differences in shape categories, supervised machine learning was employed to estimate feature importance (see Methods) [68]. Of the 67 features used as predictors in a random forest regression (see Methods), we selected only 15. OOB error is an estimate of how poorly models perform when a specific feature is excluded and is akin to error estimated from jackknife re-sampling (Fig 5). In this way, features with higher estimates tend to be more relevant for classification and prediction. In this experiment, features could only be selected up nine times, once per value of *k*. We maintained features that were selected in *≥* 3 levels of *k* to use as independent variables in classification (Table 1). The 15 selected features accounted for > 80% of importance assigned to the 67 features across all values of *k* (Fig 5B). Here, the use of “EigenFaces,” an analysis from the 1980s, designed to classify human faces, was re-purposed for the quantification and classification of fruit shape in strawberry [58, 57, 60, 59]. Pixel-based features dominated the selected features and include PCs 1 – 7 of the EigenFruit analysis (EigenFruitPC[1,7]), PCs 1 and 2 of the vertical biomass profile (BioVPC[1,2]), and PCs 1 – 3 of the horizontal biomass profile (BioHPC[1,3]) (Table 1; Fig. 5 and 6). These features originated from the same data type as used in *k*-means clustering (i.e., pixel intensities), which is likely the reason they make up the majority of the selected features (Table 1; Fig. 5 and 6). Several geometric descriptors were also selected, including the bounding aspect ratio (BAR), Shape Index (SI), and Kurtosis (Kurt) (Table 1; Fig. 5 and 6). We generated a subset of five features with mean OOB *≥* 0.069 (Fig. 5A). OOB = 0.069 was the median OOB error for all features across all classes. This subset of features included EigenFruitPC[1,2], BioVPC_1_, BioHPC_1_, and BAR (Table 1). We also generated a third smaller set that included only EigenFruitPC_1_ and BioVPC_1_ with mean OOB *≥* 0.1 (Fig. 5A). OOB = 0.1 was the mean OOB error for all features across all classes. The prevalence of pixel-based descriptors in these selected subsets indicated the magnitude of relevant shape information that they described.

**Table 1.** Broad-sense heritability of selected features

| Feature | $H^2$ | % Times Selected | Normalized Eigenvalue (80%,50%,20%) | Feature Set |
| --- | --- | --- | --- | --- |
| EigenFruit PC1 | 0.68 | 100 | 0.26 (0.27,0.27,0.26) | 15, 5, 2 |
| EigenFruit PC2 | 0.58 | 88.9 | 0.14 (0.14,0.14,0.14) | 15, 5 |
| EigenFruit PC3 | 0.00 | 55.6 | 0.05 (0.06,0.05,0.06) | 15 |
| EigenFruit PC4 | 0.69 | 55.6 | 0.04 (0.04,0.05,0.04) | 15 |
| EigenFruit PC5 | 0.43 | 66.7 | 0.03 (0.03,0.04,0.03) | 15 |
| EigenFruit PC6 | 0.47 | 55.6 | 0.03 (0.03,0.03,0.03) | 15 |
| EigenFruit PC7 | 0.29 | 44.4 | 0.02 (0.02,0.02,0.02) | 15 |
| Vertical Biomass Profile PC1 | 0.67 | 100 | 0.65 (0.66,0.66,0.66) | 15, 5, 2 |
| Vertical Biomass Profile PC2 | 0.49 | 55.6 | 0.17 (0.17,0.16,0.17) | 15 |
| Horizontal Biomass Profile PC1 | 0.65 | 88.9 | 0.44 (0.44,0.46,0.44) | 15, 5 |
| Horizontal Biomass Profile PC2 | 0.62 | 55.6 | 0.36 (0.36,0.35,0.37) | 15 |
| Horizontal Biomass Profile PC3 | 0.69 | 33.3 | 0.10 (0.10,0.10,0.10) | 15 |
| Bounding Aspect Ratio | 0.71 | 100 | NA | 15, 5 |
| Shape Index | 0.72 | 66.7 | NA | 15 |
| Kurtosis | 0.55 | 33.3 | NA | 15 |

Broad-sense heritability ( $H^2$ ) estimated on a per line basis.
% times selected is the number of classification models that a feature was selected in out of 9 (i.e., $k = [2, 10]$ ).
Normalized eigenvalues is the eigenvalue associated with a specific PC divided by the sum of all eigenvalues.
The large value is the normalized eigenvalue from the full data set. Values in parentheses contain the normalized eigenvalues for the 80%, 50%, and the 20% training sets, respectively.
Feature set indicates in which of the 3 sets a given feature was included.

**Figure 3.**
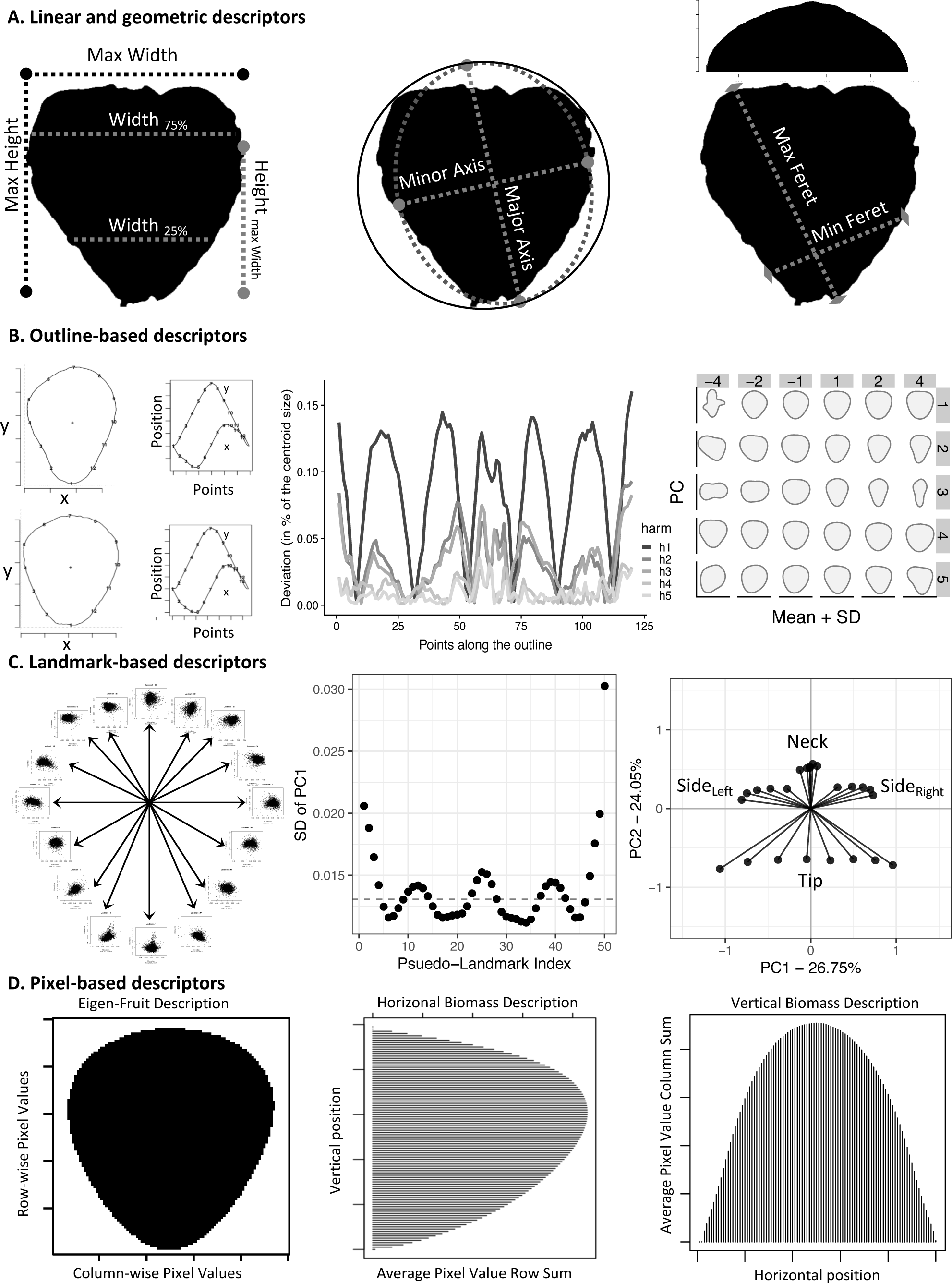
Trait Dictionary for this study. **(A)** Linear descriptors. *Left* Simple linear measurements. *Center* Best fit ellipse axes. For the circle, Round and Circ = 1. *Right* Max and Min Feret. Histogram represents the marginal distribution on the horizontal axis used to calculate Var, Skew, and Kurt. **(B)** Outline descriptors. *(Left)* The two left most images are the outlines of two strawberries with 12 evenly spaced points. The graphs on the right show the original closed outline as two oscillating functions. *(Center)* Deviations from the closed outline with increasing harmonics (harm= [*h*1, *h*5]).*(Right)* The plot shows the effects of PC [1, 5] (vertical) with effect sizes, [–4, 4] (horizontal) on the mean shape. **(C)** Landmark descriptors. *(Left)* 50 evenly spaced landmarks are extracted and treated as bi-variate features.*(Center)* Standard deviation of PC1 for each landmark is plotted in sequence. Dashed horizontal line is the median standard deviation. *(Right)* Pseudo-landmarks were selected to represent each region of high variance. Using the values on the first principal axis as observed variables, confirmatory factor analysis was performed to infer latent relationships to tip, left and right side, and neck shape. **(D)** Pixel descriptors. *(Left)* Mean EigenFruit using flattened binary images. *(Center)* Mean Horizontal Biomass using image row sums. *(Right)* Mean vertical biomass using image column sums.

**Figure 4.**
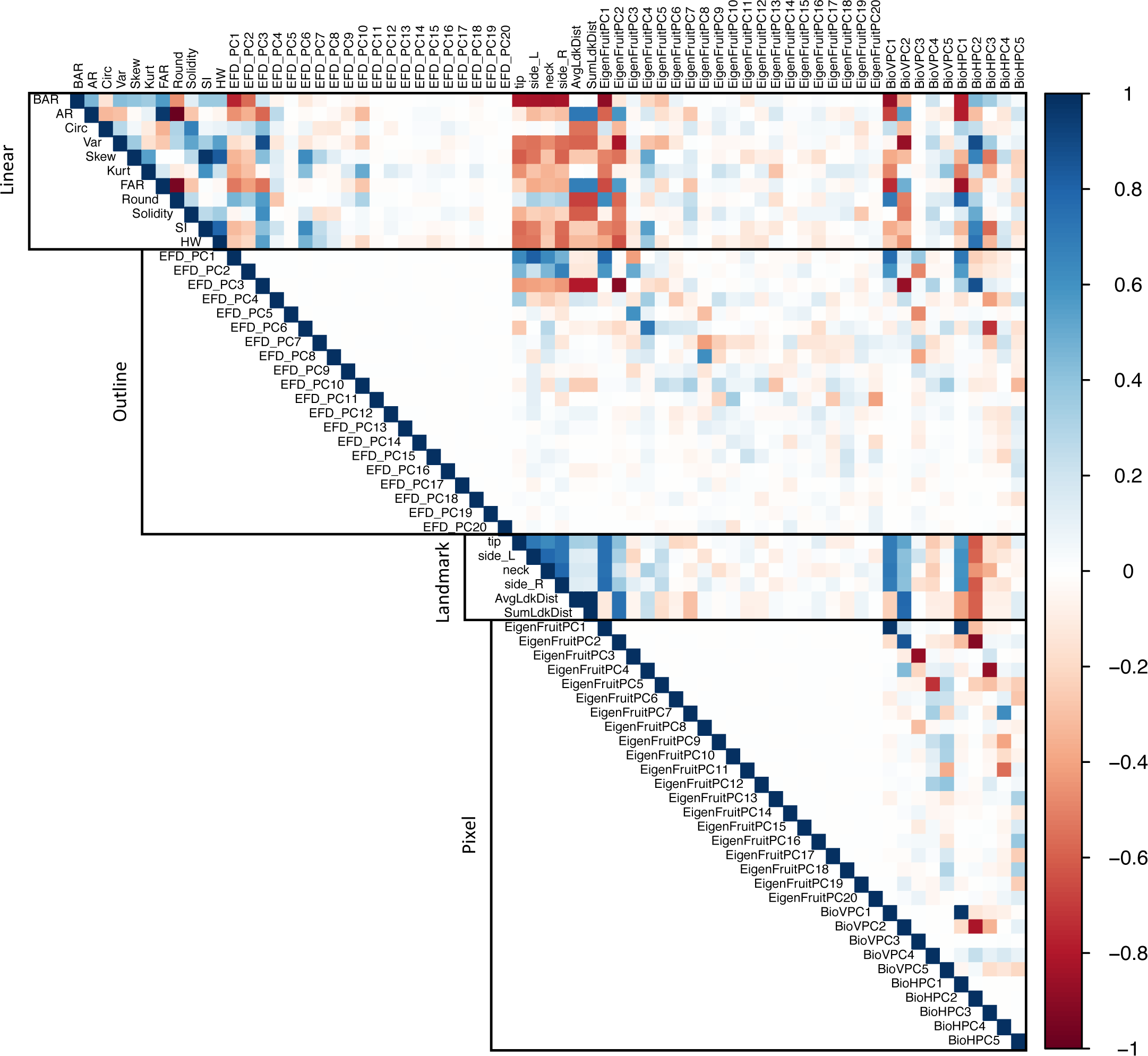
Correlations between all 67 features used in this study. Positive correlations are colored blue, negative correlations are colored red.

**Figure 5.**
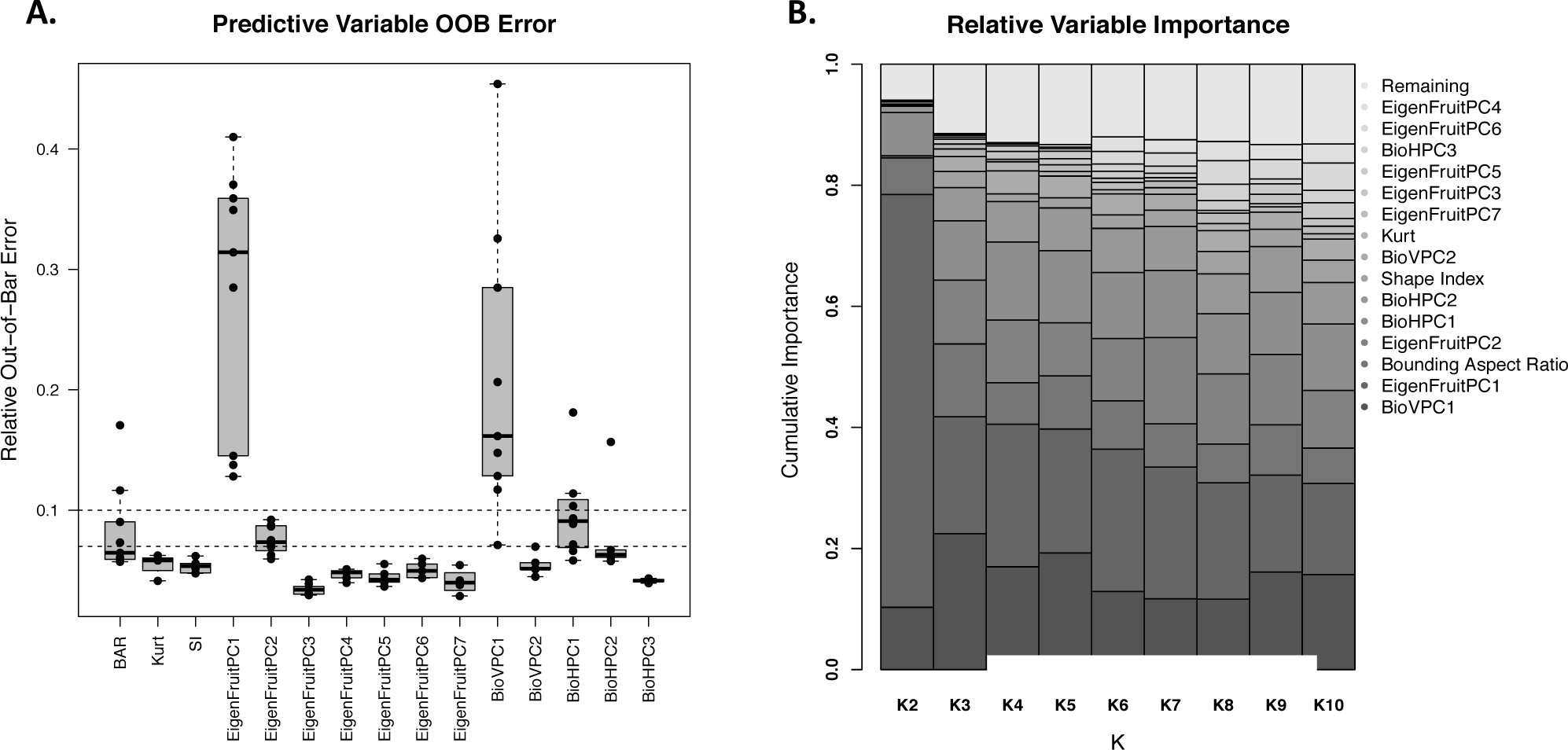
Results from feature selection. **(A)** Out-of-Bag error for each of the 15 selected features. Horizontal dashed lines are the median (0.069) and mean (0.1) OOB. **(B)** The relative importance of each feature within each level of *k*. The 15 selected features explain nearly 90% of the weight attributed to all of the features.

**Figure 6.**
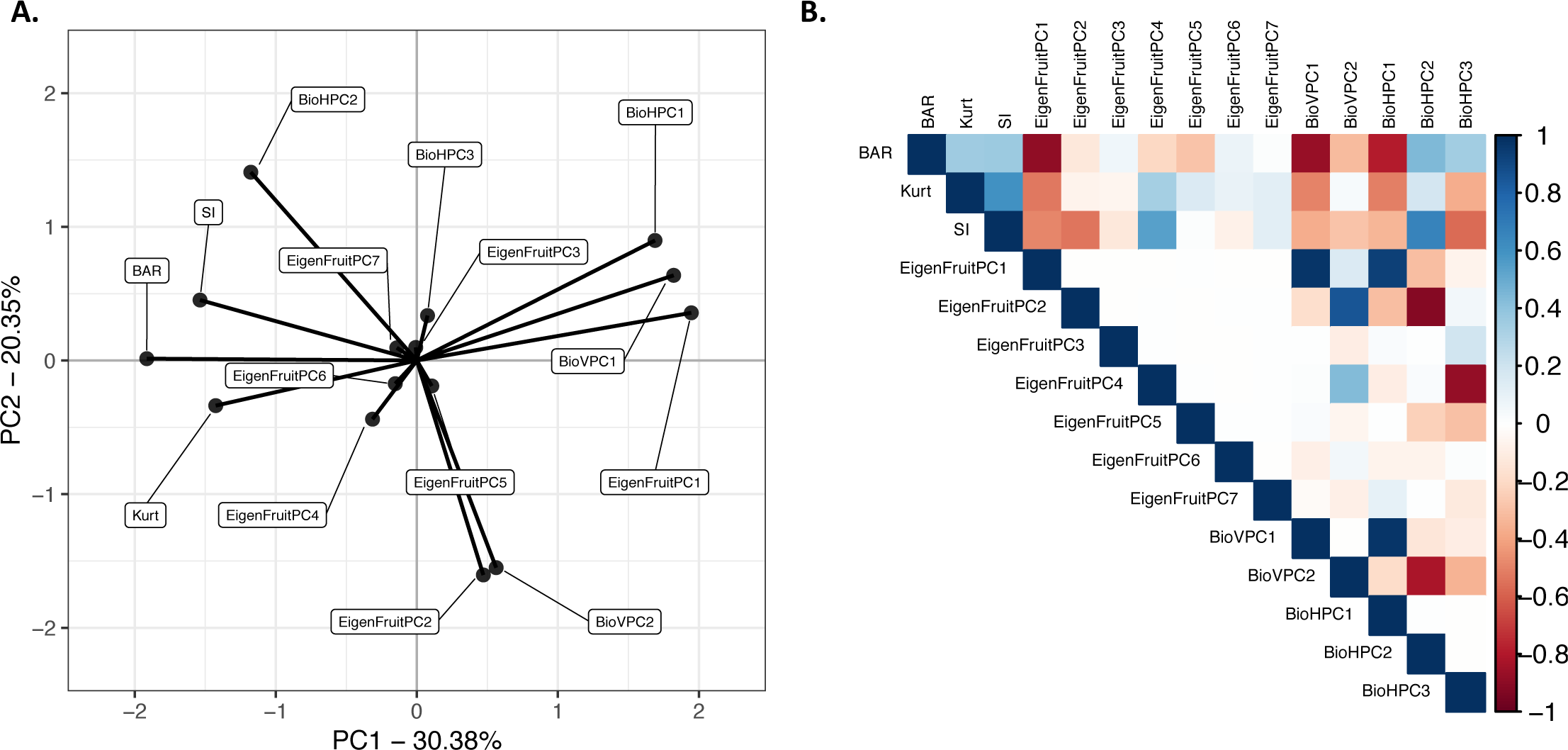
Relationship between selected features. **(A)** Principal directions of the feature variance-covariance matrix among the 15 features selected for classification. **(B)** Pearson correlation matrix of the 15 selected features. Positive correlations are colored blue, negative correlations are colored red.

### Broad-sense heritability and relationship of selected features

While the continuous nature of the morphometric features is expected to be more conducive and provide higher resolution to quantitative genetic analyses compared to their categorical counterparts, it is also vital that these features be heritable. The *H*^2^ for each feature was estimated on a clone-mean basis using a linear mixed-effect model (see Methods; Eqn. 10 and Eqn. 9) [69]. The *H*^2^ for each feature is reported in Table 1. Estimates of *H*^2^ for the quantitative features ranged from low (> 0.3) to high (> 0.7). Heritability estimates were consistent with those previously reported for shape phenotypes in strawberry and other plant species [12, 44, 50]. However, the *H*^2^ of one selected feature, EigenFruitPC_3_, was estimated to be 0.00 (Fig. S5). Similar results were reported in carrot (*Daucus carota* L.) for pixel-based root and shoot features [13] and apple (*Malus domestica*) for elliptical Fourier series leaf shape features [12]. [13] attributed the null *H*^2^ of root shape characteristics to low phenotypic variation between the inbred parents and genotype × environment interactions. While these reasons could certainly be drivers, we hypothesize that the null estimate may arise from the pixel-based descriptors describing more complex aspects of fruit or root shape. If the non-genetic component of a multi-variate phenotype is large, then performing PCA on that multi-variate trait could produce leading principal components that describe mostly non-genetic variance. However, this study and [13] are two of the only studies to report the *H*^2^ of pixel-based features in plants, and the likelihood of this phenomenon remains unclear.

Figure 6A shows the directions of the feature variance-covariance matrix with the traits labeled as in Figure 5. Figure 6B shows the correlation matrix between the 15 selected features. For the five features selected by OOB error (Fig. 5), indicated with a 5 in Table 1, the estimated *H*^2^ was *≥* 0.58. As the majority of selected features are principal components of different pixel-based analyses (Fig. S6), there were many weak correlations (Fig. 6B). We hypothesize that the importance of these features is partly driven by the similarity of the raw data (i.e., binary pixel intensities) used in *k*-means clustering to acquire shape categories and for EigenFruit shape analysis. Although principal components are uncorrelated, we observed strong correlations between PCs from different analyses (Fig. 6). EigenFruitPC_1_ shared a strong positive correlation with both BioVPC_1_ and BioHPC_1_ (ρ = 0.98; *p* < 2*e* – 16 and ρ = 0.93; *p* < 2*e* – 16, respectively), as did EigenFruitPC_2_ with BioVPC_2_ (ρ = 0.86; *p* < 2*e* – 16). BioHPC_2_ was negatively correlated with both EigenFruitPC_2_ and BioVPC_2_ (ρ = –0.92; *p* < 2*e* – 16 and ρ = –0.81; *p* < 2*e* –16, respectively). BioHPC_3_ was negatively correlated with EigenFruitPC_4_ (ρ = –0.87; *p* < 2*e* –16). BAR was negatively correlated with EigenFruitPC_1_, BioVPC_1_ and BioHPC_1_ (ρ = –0.89; *p* < 2*e* –16, ρ = –0.87; *p* < 2*e* –16, and ρ = –0.78; *p* < 2*e* –16, respectively). The correlations between these features indicated that the pixel-based descriptors describe comparable patterns of phenotypic variation.

### Image Classification using Selected Features

The accuracy of classification, or prediction, is typically assessed by cross-validation [22, 70]. We generated training sets that consisted of 80%(5, 500), 50% (3, 437), or 20% (1, 374) of the images. Assignment to either training or test set was random and without stratification. *k*-means clustering was performed using the training sets, and *k* was allowed to range from 2 to 10. We assigned the test set images to the nearest neighboring cluster for each level of *k*. We performed PPKC on the clusters derived from the training set and the similarity between the full set and training sets were visually assessed. The clusters derived from the different sets appeared to be nearly identical (Fig. S7). The order of clusters derived from the reduced data set also appears identical to those described in the full set (Fig. S7). The principal component-based features were recalculated using the training data sets and the corresponding test set images projected into the new space. We only extracted the 15 selected features. These included EigenFruitPC[1,7], BioVPC[1,2], and BioHPC[1,3] (Table 1). The selected geometric features, including BAR, SI, and Kurt, were not recalculated as they do not change concerning the other samples, unlike *k*-means and PCA which both rely on and change based on observed data. For EigenFruitPC[1,7], BioVPC[1,2], and BioHPC[1,3], the percent variance explained by each feature was similar to that in full data set (Table 1), indicating that the principal components derived from the reduced set describe similar features of shape as those derived from the full set.

Support vector regression (SVR) and linear discriminant analysis (LDA) were both used for classification (see Methods). We performed ten iterations of each set size and feature set across all levels of *k*. The results of this experiment are reported in Table 2. Overall, the models performed with high accuracy of classification. Generally, as we used fewer features for classification model performance is reduced, most notably for larger values of k. Indeed, when *k* = 2 accuracy improved slightly with fewer features in the different models. Except for one case, SVR was found to outperform linear discriminant analysis consistently. The one case where LDA outperformed SVR was when *k* = 10, including two features, and 20% training data. LDA achieved 69.4% accuracy, and SVR achieved 68.7% accuracy. Using five features for classification, we achieve the highest accuracy (99.3%) for *k* = 2. In the range of interest, *k* = [2, 4], the models do not fall below 94.0% accuracy for any training set size.

**Table 2.**
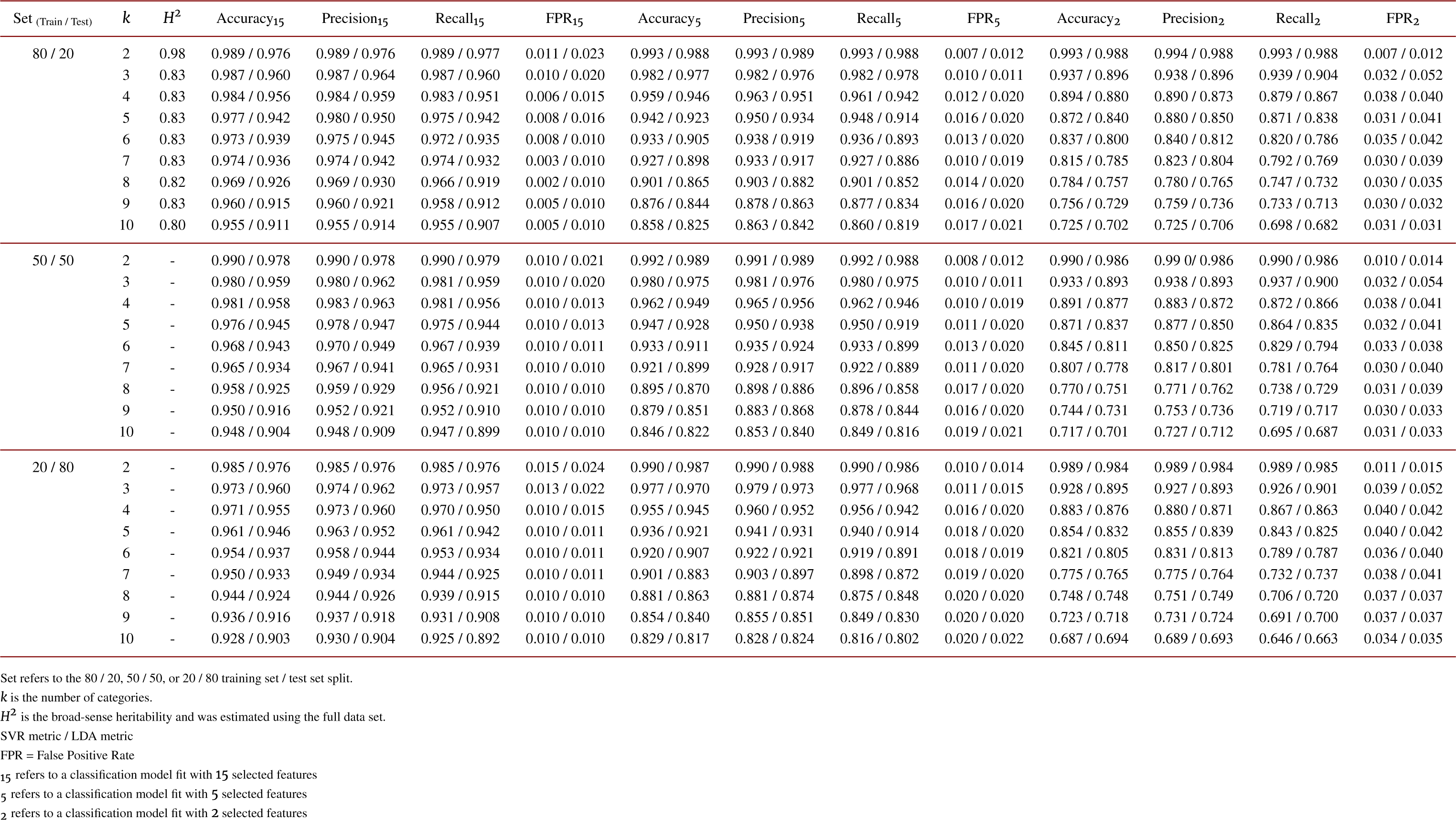
Classification model evaluations validation experiment

## Discussion

As high-throughput phenotyping for external fruit characteristics becomes more and more widely of interest to specialty crop researchers, we expect that this work will have various applications in both applied and basic plant research [49, 12, 13], cultivar development [14, 44], intellectual property protection and documentation [71, 72], and waste reduction [18, 73]. Our study showed that strawberry fruit shapes could be robustly quantified and accurately classified from digital images. Most importantly, our analyses yielded quantitative phenotypic variables that describe fruit shape (Fig. 3), arise from continuous distributions, and are moderately to highly heritable (*H*^2^) (Table 1) [35, 36]. We accomplished this by translating two-dimensional digital images of fruit into categorical and continuous phenotypic variables using unsupervised machine learning and morphometrics, respectively [12, 13, 49, 58, 60, 74]. We found that mathematical approaches developed for human-face recognition [57, 58] were powerful for strawberry fruit shape recognition (Table 1), that unsupervised shape clustering was robust to sample size deviations (Fig. S7), and that only a few quantitative features are needed to accurately classify shapes from images (Table 2).

We empirically derived the shape progression produced in the present study through the application of PPKC (Alg. 1; Fig. 2). Ordinal categorical traits are commonplace in quantitative genetic studies [27, 75] and are the current standard for phenotyping external fruit characteristics [14, 15, 44]. PPKC identified four exemplary strawberry shape categories in the population we studied, which were characterized by a progression from ‘longer-than-wide’ (prolate) to ‘wider-than-long’ (oblate) (Fig. 2). Critically, this gradient agreed with the arbitrarily defined progressions in previous reports [14, 16]. However, unlike previous studies, which suggested using nine [14] or eleven shape categories [18], our work presented empirical evidence for a specific number of mathematically defined shape categories. We determined that *k* = 4 was the appropriate level of complexity based on the visual appearance of the discovered clusters (Fig. 2), high *H*^2^ estimates (Table 2), and the information criteria calculated for the *k*-means models (Fig. S3). Interestingly, PPKC can determine a visually, reasonable phenotypic gradient up to *k* = 7 (Fig. S4) despite strong evidence of overfitting for *k* > 4 (Fig. S3). Because unsupervised clustering does not define unobserved categories, more shape categories may exist mainly between individuals with more extreme or more variable fruit shape phenotypes or greater genetic distances.

The specific genetic factors that give rise to variation in fruit shape in garden strawberry are currently unknown and have been understudied in strawberry [44]. The selective pressure exerted on fruit shape in strawberry could have impacted large-effect loci, in which case ordinal phenotypic scores are likely to be sufficient for identifying genetic factors affecting fruit shape. Loss-and gain-of-function mutations have played an essential role in identifying genes affecting fruit shape in tomato, a model that has been highly instructive and important for understanding the genetics of fruit shape and enlargement in plants [32, 33, 38, 76, 77]. There are striking examples in tomato and other plants where identified genes regulate the development of fruit shape. For example, the *OVATE* gene in tomato regulates the phenotypic transition from round- to pear-shaped fruit [78, 79]. If large-effect mutations underlie differences in strawberry fruit shape, the ordinal classification system proposed here should enable the discovery of such effects. Furthermore, quantitative phenotypes were linked to genetic features that interact with large-effect genes, i.e., suppressors of *OVATE (sov)*, through bulk segregant analysis and QTL mapping [80]. In woodland strawberry (*F. vesca*), fruit size and shape are linked to the accumulation and complex interaction of auxin, GA, and ABA, mediated by the expression and activity of *FveCYP707* and *FveNCED*, as well as other genes [9]. Because of the high *H*^2^ estimates for several of the newly created phenotypic variables 1, we hypothesize that quantitative shape phenotypes can yield a more comprehensive understanding of the underlying genetic mechanisms of fruit shape in garden strawberry (*F*. × *ananassa*) through genome-wide association studies and other forward-genetic analyses [35, 36, 81]. We anticipate that the analyses in this study will enable us to discover and study the genetic determinants of fruit shape in strawberry and other specialty crops.

## Methods

### Mating and Field Design

Seventy-five bi-parental crosses were generated by controlled pollination of 30 parents in an incomplete (16 × 14) factorial mating design. These parents were chosen to represent a broad range of phenotypic diversity in the University of California, Davis strawberry germplasm. 2, 800 hybrid progeny were planted at the Wolfskill Experimental Orchard in Winters, CA in sets of 20 or 40 per family, depending on seedling survival. 20% of the planted materials from each family were randomly selected for further testing. Clones of 545 of the selected 560 progeny were successfully propagated. 12 bare-root runner plants of each of the 545 progeny and the 30 parents were collected and planted in November 2017 in Salinas, CA in 4 plant plots as a randomized complete block design with three replicates of each genotype.

### Image acquisition

Strawberries were harvested from plots in Salinas, CA once in April 2018 and again in May 2018. Digital images of up to 3 fruit per plot were imaged using a Sony α-6000 Mirrorless digital camera mounted on a portable copy stand in aperture priority, with the aperture set to f/8. Strawberries with the calyx removed were placed in the frame against a black felt backdrop, along with a QR-code identifying the plot, such that the most extensive face was perpendicular to the sensor. Berries were mounted to set of staples to eliminate any rolling or pitch of the berries. The time to stage a given set of fruit and acquire an image ranged from 1 to 2 min. All images were acquired with a 16 – 50 mm lens set to 16 mm and positioned approximately 16 cm above the base of the copy stand resulting in images with 97.4 pixels per cm. In total, 2, 924 plots were imaged over the two harvest dates.

### Image Processing

Input files were JPEG images (3008px*×*1688px) with the strawberries placed in regular positions within a scene. All images were first segmented and converted to binary using the Simple Interactive Object Extraction (SIOX) tool in ImageJ 2.0.0 [82, 83, 84] through custom batch scripts. Images that were unsuccessfully segmented were flagged and handled individually to ensure completeness. ImageJ was used to acquire the bounding rectangle of each object of interest. Each object was extracted based on the dimensions of its bounding rectangle using R 3.5.3 [85] and the jpeg package [86]. White pixels were added to the edges of each image such that the resulting images is a square of size max(*H, W*) × max(*H, W*) using the “magick::image_border()” package [87]. “magick::image_resize()” was used to scale the images from max(*H, W*) × max(*H, W*) px to 1000 × 1000 px. This method results in binary images that maintain the original aspect ratio with a maximum dimension equal to 1, 000 pixels and then resized to 100 × 100.(Fig. 1). In total, the downstream analyses included 6, 874 images of individual berries.

### Feature extraction

#### Categorical features

This method afforded clustering decisions based on raw image data instead of the extracted quantitative features. Each image matrix was flattened into a single 10, 000 element row vector; all of the samples were then bound together by columns. The resulting matrix for all samples was 6, 874 × 10, 000. The “stats::kmeans()” function in R was used to perform *k*-means clustering. Values of *k* (i.e., the number of clusters) range from 2 to 10. Assigned clusters were recorded for all values of *k*.

After initial clustering, the centroids of each cluster from each value of *k* were visually examined. We determined that, when *k ≥* 8, there were two (*≥* 2) groups that appeared to be mirror images of each other. Mirrored groups likely arose as artifacts of the imaging set up and symmetries between perspectives and are likely not reflective of any true biological characteristic. All images in one of the mirrored groups were reloaded, rotated around their vertical axis, and flattened into vectors as before. The “stats::kmeans()” function clustered this modified data set allowing *k* to vary between 2 and 10. We recorded newly assigned clusters each object for all values of *k*. A second visual inspection concluded that the modification removed mirror groups when *k ≥* 4. We used the modified clusters assignments in downstream analyses. Modified clusters are then reordered using PPKC (Fig. 2). The ordered categories, across the various levels of *k*, became the response for classification experiments. The correct choice of k is often ambiguous, with interpretations depending on the shape and scale of the distribution of points in a data set and the desired clustering resolution of the user. In addition, increasing k without penalty will always reduce the amount of error in the resulting clustering, to the extreme case of zero error if each data point is considered its own cluster (i.e., when k equals the number of data points, n). Intuitively then, the optimal choice of k will strike a balance between maximum compression of the data using a single cluster, and maximum accuracy by assigning each data point to its own cluster. The optimal value of *k* was determined based on four different evaluation criteria: total within-cluster sum of squares, adjusted R2, AIC, and BIC.

#### Linear and geometric features

Linear and geometric features measure aspects of the fruit directly from images and were processed using ImageJ 2.0.0 [83, 84] and R 3.5.3 [85]. Extracted measurements included Shape Index (SI) [42], Circularity (Circ) [84], Bounding Aspect Ratio (BAR) [84], Ellipse Aspect Ratio (AR) [84], Roundness (Round) [84], Solidity (Solid) [84], Feret Aspect Ratio (FAR) [84], the ratio of the height of max width and max height (HW) [42], Variance (Var), Skewness (Skew) [84], and Kurtosis (Kurt) [84] (Fig. 3A). For Var, Skew, and Kurt, the analyses focus on the horizontal axis (Fig. 3A).

#### Elliptical Fourier features

EFA comprehensively described closed outlines as a series of oscillating, harmonic functions and were calculated using Momocs v1.2.9 [88] in R 3.5.3. We extracted elliptical Fourier features for the first 5 harmonics resulting in 20 coefficients using “Momocs::efourier()” function. Each harmonic level is made up of 4 coefficients that correspond to the effects of the cosine and sine in the *x*-axis (coefficients *A* and *B*) and the *y*-axis (coefficients *C* and *D*). To allow for discrimination between accessions based on fruit shape, principal component analysis (PCA) was performed using the “Momocs::PCA” from Momocs for EFFs. We recorded the eigenvectors of each image on the 20 resulting principal axes (Fig. 3B).

#### Generalized Procrustes and revealed latent features

GPA describes the shape as the distance either between landmarks and a centroid. The outline of each object was decomposed into 50 evenly spaced pseudo-landmarks moving clockwise around the object. The “Momocs::fgProcrustes()” function from Momocs v1.2.9 [88] was used to perform the alignment between shapes (Fig. 3C; left). Each of the 50 aligned pseudo-landmarks was considered as an individual multi-variate feature. Each of the 50 features was centered such that the marginal mean of both axes is 0. The “stats::prcomp()” function in R was used to perform PCA on each of the 50 centered pseudo-landmarks (Fig. 3C; left and center).

Latent features from the calculated landmark principal components were constructed to describe the 4 most variable regions of the strawberry outline (i.e., tip, left side, neck, and right side) (Fig. 3C; center) with “lavaan::sem()” using the lavaan package *v*0.6–3 [89]. Use of SEM is commonly justified in the social sciences because of its ability to impute relationships between unobserved constructs (latent variables) from observable variables. Here, we treat different the pseudo-landmarks as the observable variables to impute the relationship between latent components of shape. Only those pseudo-landmarks with variance on PC1 greater than the median were used to manifest the four latent features (Fig. 3C; center and right). Latent features were established by the following structure:

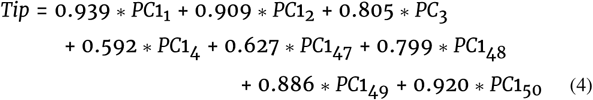

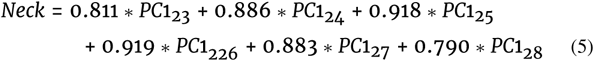

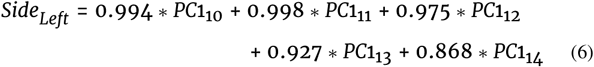

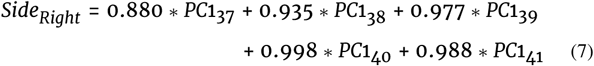

The weights for each variable are those reported from the “lavaan::sem()” function. *Tip* is manifested by a combination of PC1 of the pseudo-landmarks 1, 2, 3, 4, 47, 48, 49, and 50 (4); *Neck* by PC1 of landmarks 23, 24, 25, 26, 27, and 28 (5); *Side*_*Left*_ by PC1 of landmarks 10, 11, 12, 13, and 14 (6); and *SideRight* by PC1 of landmarks 37, 38, 39, 40, and 41 (7). Each of the four latent features were calculated for all images. The fit of this model was determined to be adequate based on the *SRMR* = 0.092, *RMSEA* = 0.068 *±* 0.001, and *CFI*/*TFI* = 0.981/0.978 [90].

#### EigenFruit features

EigenFruit features were adapted from the EigenFaces methods of [57] and [58] and incorporate information about every pixel in an image. The resulting matrix of binary image vectors was 6874 × 10, 000. There can only be as many non-zero PC’s as there are observations (i.e., 6, 874). The “stats::prcomp()” function was used to perform PCA. We recorded the eigenvalues of the first 20 PCs. Together these 20 PCs explained 71.7% of the variance. PC1, PC2, and PC3 explained 26.8%, 12.6%, and 5.24%, respectively (Fig. 3D; left).

#### Biomass profile features

Biomass profile features describe the shape as the sum of pixels in each row, or column, of a given image. We adopted this method from [13]. We generated the horizontal biomass profile by recording the number of black pixels in each of 100 rows. The vertical biomass profile was generated by recording the number of black pixels in each of the 100 columns. The function “stats::prcomp()” in R was used to perform PCA for each profile (i.e., vertical and horizontal). The eigenvectors of the first 5 PCs from each were retained. Together these 5 PCs explained 95.9% and 95.4% of the total symmetric shape variance for the horizontal and vertical profiles, respectively (Fig. 3D; center and right).

### Broad-sense Heritability Estimation

#### Qualitative Features

Broad-sense heritability on a clone-mean basis (*H*^2^) for each ordered level of *k* was estimated using the ordinal package v2019.3 – 9 [67] in R 3.5.3. Variance components were estimated using a cumulative link mixed models with a cumulative logit link function and a multinomial error,

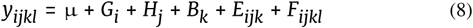

*y*_*ijkl*_ is the categorical feature, μ is the grand mean, *G*_*i*_ is the random effect of *i*th genotype 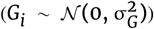, *H*_*j*_ is the fixed effect of the *j*th harvest, *B*_*k*_ is the fixed effect of the *k*th block, *E*_*ijk*_ is the residual error of the *ijk*th plot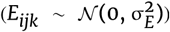, and *F*_*ijkl*_ is the error of *ijkl*th fruit (subsample) 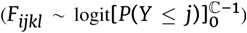. The “clmm()” function implements of cumulative link mixed models for ordinal data. Ordinal GLMMs were considered the most appropriate, and conservative, approach because we could not assume that shape categories would be linear. Variance component estimation is performed via maximum likelihood and allows for multiple random effects with crossed and nested structures [67]. *H*^2^ for each feature was calculated as

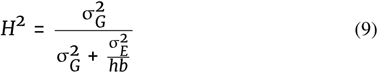

Where 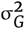 is the genetic variance, 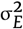 is the residual variance, *h* is the harmonic mean of observed harvest dates per genotype (1.94), and *b* is the harmonic mean of observed blocks per genotype (2.89).

#### Quantitative Features

Broad-sense heritability on a clone-mean basis (*H*^2^) was estimated for features with the lme4 package v1.1 – 19 [69] in R 3.5.3. REML variance components were estimated using the linear mixed effect model,

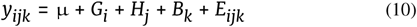

*y*_*ijk*_ is the quantitative feature, μ is the grand mean, *G*_*i*_ is the random effect of the *i*th genotype 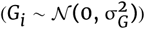, *H*_*j*_ is the fixed effect of the *j*th harvest, *B*_*k*_ is the fixed effect of the *k*th block, *E*_*ijk*_ is the residual error of the *ijk*th plot 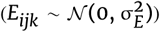. Only two Harvest dates and three Blocks were observed and, because of this, they were treated as fixed effects. *H*^2^ for each feature was calculated as in equation 9. Where 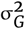 is the genetic variance, 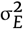 is the residual variance, *h* is the harmonic mean of observed harvest dates per genotype (1.94), and *b* is the harmonic mean of observed blocks per genotype (2.89).

### Feature selection

Random forest regression models were fit in R 3.5.3 using the VSURF package v1.0.4 [68]. 100 forests, each consisting of 2, 000 random trees were fit using 67 features to predict cluster assignments. The “VSURF::VSURF()” function returns two sets of features. The first includes important features with some redundancy, and the second, smaller set, corresponds to a model focusing more closely on the classification and reducing redundancy [68]. Features that appeared in the second set for more than three levels of *k* were recorded and used for classification for all clusters (feature set 15). Five features had mean OOB estimates greater than the median (OOB= 0.069) were used as feature set 5. Two features had mean OOB estimates greater than the mean estimate (OOB= 0.1) were recorded as feature set 2.

### Classification performance

The classification accuracy was then estimated using the “MASS::lda()” function from MASS v7.3 – 51.1 [91] as well the “e1071::svm()” function from e1071 v1.7–0 [92]. Classification models were trained to delineate the cluster assignments from modified *k*-means using the three different feature sets as predictor variables. All images were randomly sorted into training and test sets without stratification of size 80/20%, 50/50%, and 20/80% to explore the relationship between sample size and model performance. The training set images were clustered using the “stats::kmeans()” function in R. As before, *k* was allowed to range from 2 to 10 for this experiment. The images in the test set were assigned to the nearest cluster for each value of *k*. The principal component features (i.e., EigenFruitPC[1, 7], BioVPC[1, 2], and BioHPC[1, 3]) were calculated using only the training set images and the test images were projected into this new space. The maximum number of non-zero principal components in this experiment for the EigenFruit analysis was either 5, 500, 3, 437, or 1, 374, depending on the size of the training data set. The PVE of each leading PC was recalculated. Geometric descriptors (i.e., BAR, SI, and Kurt) were not recalculated as they are derived from an individual sample and not a sample population. Finally, both LDA and SVR models were trained using all three feature sets for all values of *k* using the “MASS::lda()” and “e1071::svm()” functions in R. The trained models were used to classify the images in the respective test set. The model performance was evaluated using the average classification accuracy, precision, recall, and false positive rate (FPR) of 10 iterations of cross validation.

## Availability of source code and requirements

Lists the following:

- Project name: 2DShapeDescription
- Project home page:https://github.com/mjfeldmann/2DShapeDescription
- Operating system(s): Platform independent
- Programming language: R and ImageJ Macro
- Other requirements: Not Applicable
- License: MIT License.
- Any restriction to use by non-academics: none

## Availability of supporting data and materials

The data supporting the results of this article are available in the Zenodo repository [63]. The code to reproduce these analyses are documented and available on GitHub [64].

## Additional files

The additional files for this article are available in the Zenodo repository [63].

- **Additional file 1: Fig. S1** Modified *k*-means clustering. (A) Results of *k-*means clustering performed on flattened binary images. (B) (1) Resulting centroids are visualized and inspected for abnormalities. In this example, two of the 8 classes, 2 and 6, appear to be mirror images. (B) (2) All images in second class are rotated on the vertical axis. (B) (3) Similarity is visually inspected by overlaying the rotated centroid onto the other, non-rotated centroid. In this example, the overlay exposes a high level of reflective symmetry. (C) *k*-means clustering is performed again for all levels of *k* but with all images assigned to class 6 rotated on the vertical axis. The lines representing each clusters centroid reflect the 20th, 40th, 60th, and 80th quantiles, moving out from the center of each images.
- **Additional file 2: Fig. S2** Results of PPKC against original cluster assignments. Ordered centroids from *k* = 2 to *k* = 8. On the left are the unordered assignments from *k*-means, and the on the right are the order assignments following PPKC. Cluster position indicated on the right [1, 8].
- **Additional file 3: Fig. S3** Optimal Value of *k*. (A) Total within cluster sum of squares. (B) Inverse of the Adjusted R2. (C) Akaike information criterion (AIC). (D) Bayesian information criterion (AIC). All metrics were calculated on a random sample of 3, 437 images (50%). 10 samples were randomly drawn. The vertical dashed line in each plot represents the optimal value of *k*. Reported metrics are standardized to be between [0, 1].
- **Additional file 4: Fig. S4** Hierarchical clustering and distance between classes on PC1. The relationship between clusters at each value of *k* is represented as both a dendrogram and as bar plot. The labels on the dendrogram (i.e., V1, V2, V3,…, V10) represent the original cluster assignment from *k*-means. The barplot to the right of each dendrogram depicts the elements of the eigenvector associated with the largest eigenvalue form PPKC. The labels above each line represent the original cluster assignment.
- **Additional file 5: Fig. S5** BLUPs for 15 selected features. For each plot, the X-axis is the index and the Y-axis is the BLUP value estimated from a linear mixed model. Grey points represent the mean feature value for each individual. Each point is the BLUP for a single genotype.
- **Additional file 6: Fig. S6** Effects of Eigenfruit, Vertical Biomass, and Horizontal Biomass Analyses. (A) Effects of PC [1, 7] from the Eigenfruit analysis on the mean shape (center column). Left column is the mean shape minus 1.5× the standard deviation. Right is the mean shape plus 1.5× the standard deviation. The horizontal axis is the horizontal pixel position. The vertical axis is the vertical pixel position. (B) Effects of PC [1, 3] from the Horizontal Biomass analysis on the mean shape (center column). Left column is the mean shape minus 1.5× the standard deviation. Right is the mean shape plus 1.5× the standard deviation. The horizontal axis is the vertical position from the image (height). The vertical axis is the number of activated pixels (RowSum) at the given veritcal position. (C) Effects of PC [1, 3] from the Vertical Biomass analysis on the mean shape (center column). Left column is the mean shape minus 1.5× the standard deviation. Right is the mean shape plus 1.5× the standard deviation. The horizontal axis is the horizontal position from the image (width). The vertical axis is the number of activated pixels (ColSum) at the given horizontal position.
- **Additional file 7: Fig. S7** PPKC with variable sample size. Ordered centroids from *k* = 2 to *k* = 5 using different image sets for clustering. For all *k* = [2, 5], *k*-means clustering was performed using either 100, 80, 50%, or 20% of the total number of images; 6, 874, 5, 500, 3, 437, and 1, 374 respectively. Cluster position indicated on the right [1, 5].

## Declarations

EFA: Elliptical Fourier Analysis; PPKC: Principal Progression of K Clusters; GM: Geometric Morphometrics; GPA: Generalized Procrustes Analysis; OOB: Out-of-Bag Error; PC: Principal Component; FPR: False Positive Rate; SVR: Support Vector Regression; LDA: Linear Discriminant Analysis; PVE: Percent Variance Explained; SIOX: Simple Interactive Object Extraction; VSURF: Variable Selection Using Random Forests.

## Competing Interests

The author(s) declare that they have no competing interests.

## Funding

This research was supported by grants to S.J.K. from the United Stated Department of Agriculture (http://dx.doi.org/10.13039/100000199) National Institute of Food and Agriculture (NIFA) Specialty Crops Research Initiative (# 2017-51181-26833) and California Strawberry Commission (http://dx.doi.org/10.13039/100006760), in addition to funding from the University of California, Davis (http://dx.doi.org/10.13039/100007707).

## Author’s Contributions

The overall project was conceived by MJF and SJK; MAH, RAF, CML, and GSC helped grow plant material and collect raw data; MJF performed the analyses; MJF, AT, and SJK wrote the paper.

## Acknowledgements

We thank Bruce Campopiano and Eduardo Garcia for assistance with several aspects of the field experiments. We thank Patrick J. Brown, Daniel H. Chitwood, Christine H. Diepenbrock, Sarah D. Turner, and Daniel E. Runcie for their comments and advice and for reviewing this manuscript.

Opinions, findings, conclusions, or recommendations expressed in this publication are those of the authors and do not necessarily reflect the views of the USDA. USDA is an equal opportunity provider and employer.

---

**Figure S1.**
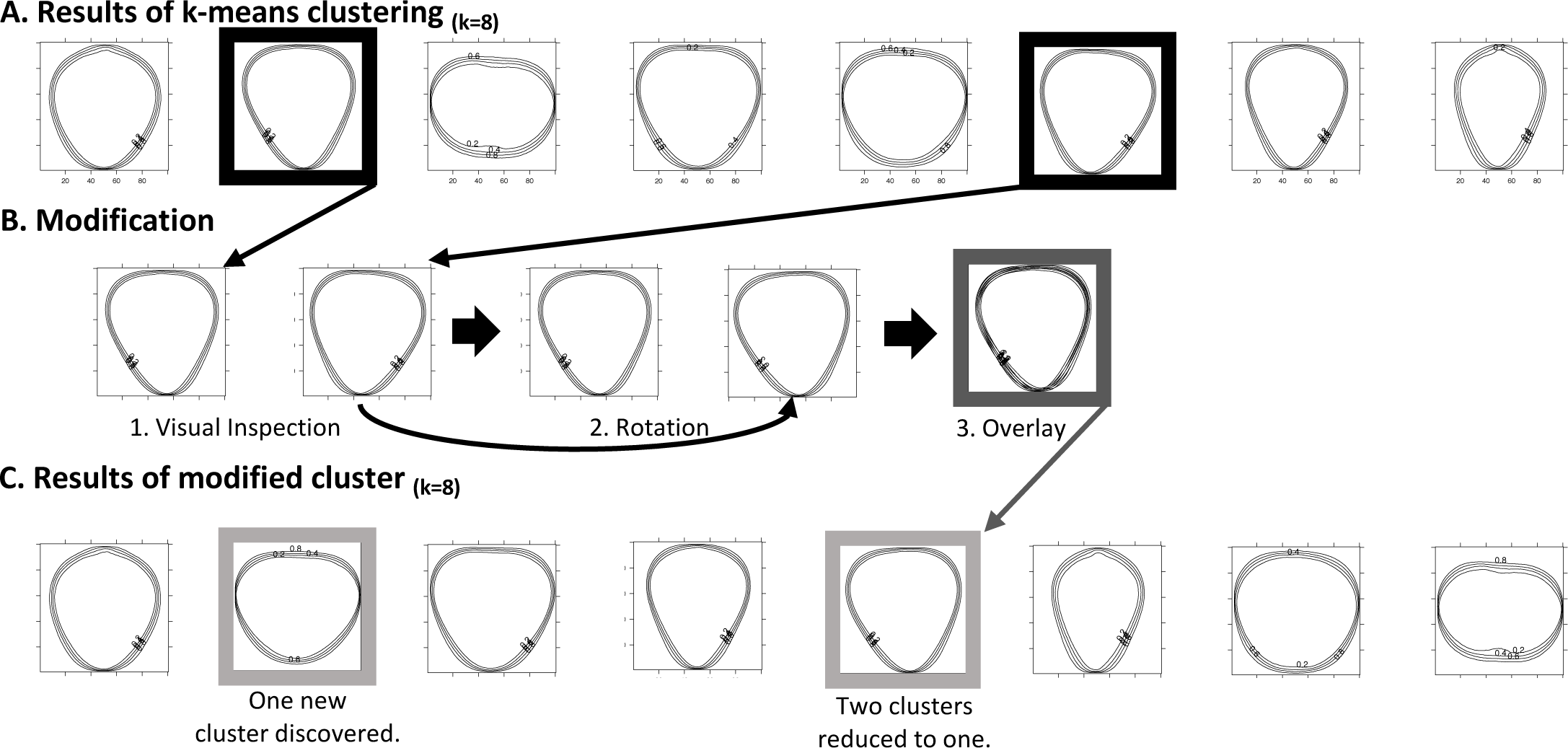
Modified *k*-means clustering. **(A)** Results of *k*-means clustering performed on flattened binary images. **(B)** (1) Resulting centroids are visualized and inspected for abnormalities. In this example, two of the 8 classes, 2 and 6, appear to be mirror images. **(B)** (2) All images in second class are rotated on the vertical axis. **(B)** (3) Similarity is visually inspected by overlaying the rotated centroid onto the other, non-rotated centroid. In this example, the overlay exposes a high level of reflective symmetry. **(C)** *k*-means clustering is performed again for all levels of *k* but with all images assigned to class 6 rotated on the vertical axis. The lines representing each clusters centroid reflect the 20th, 40th, 60th, and 80th quantiles, moving out from the center of each images.

**Figure S2.**
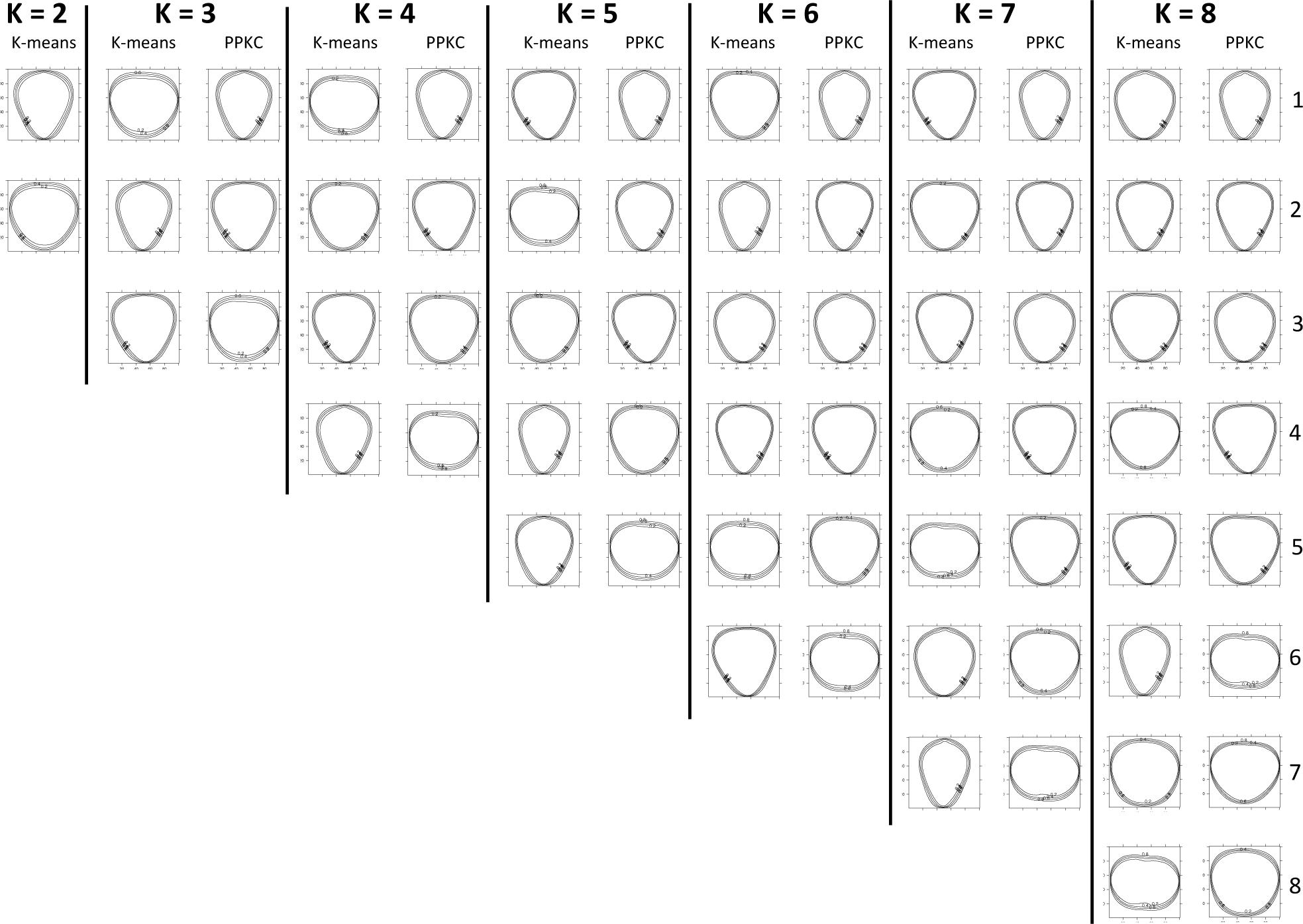
Results of PPKC against original cluster assignments. Ordered centroids from *k* = 2 to *k* = 8. On the left are the unordered assignments from *k*-means, and the on the right are the order assignments following PPKC. Cluster position indicated on the right [1, 8].

**Figure S3.**
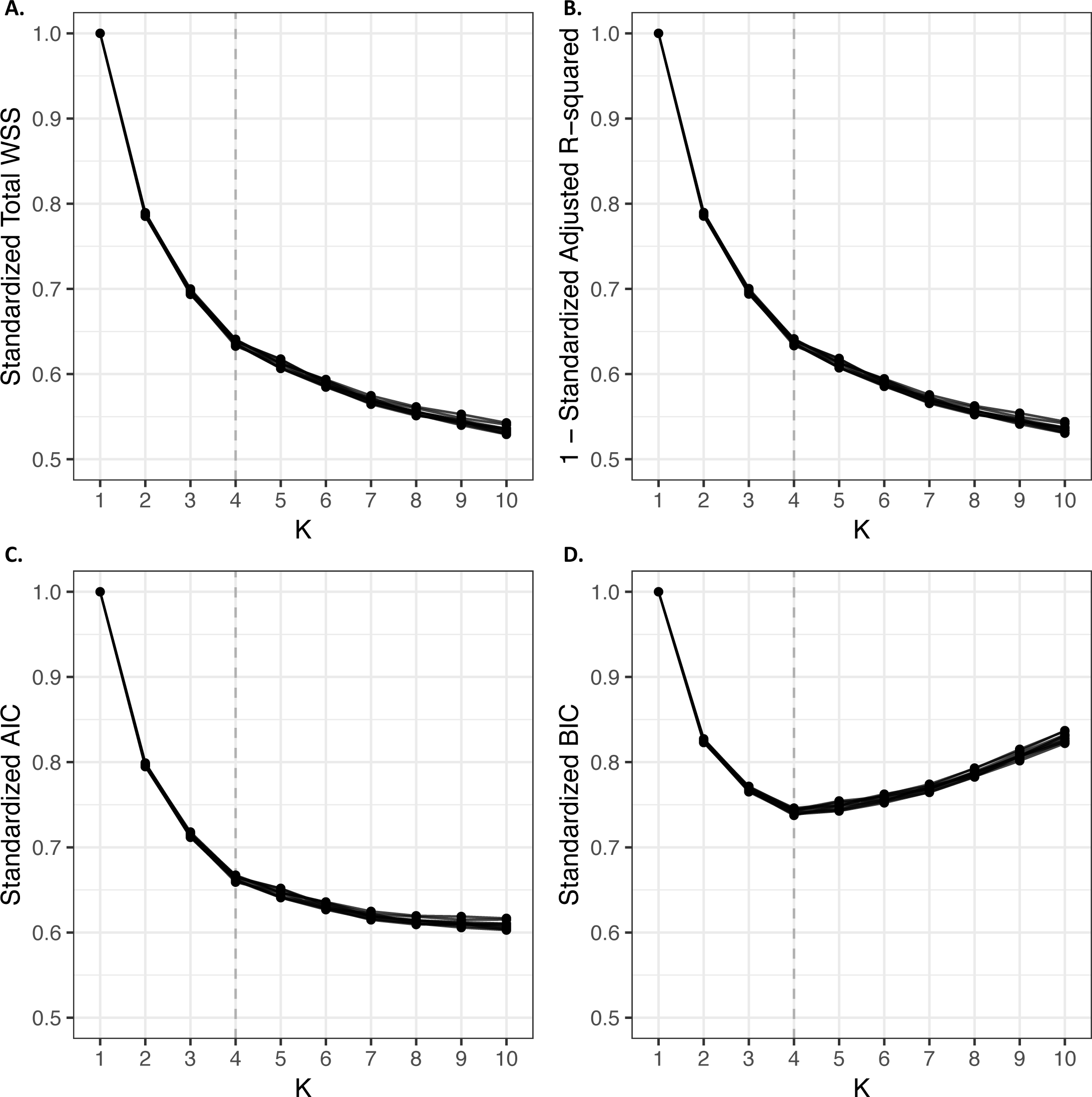
Optimal Value of *k*. **(A)** Total within cluster sum of squares. **(B)** Inverse of the Adjusted R^2^. **(C)** Akaike information criterion (AIC). **(D)** Bayesian information criterion (AIC). All metrics were calculated on a random sample of 3, 437 images (50%). 10 samples were randomly drawn. The vertical dashed line in each plot represents the optimal value of *k*. Reported metrics are standardized to be between [0, 1].

**Figure S4.**
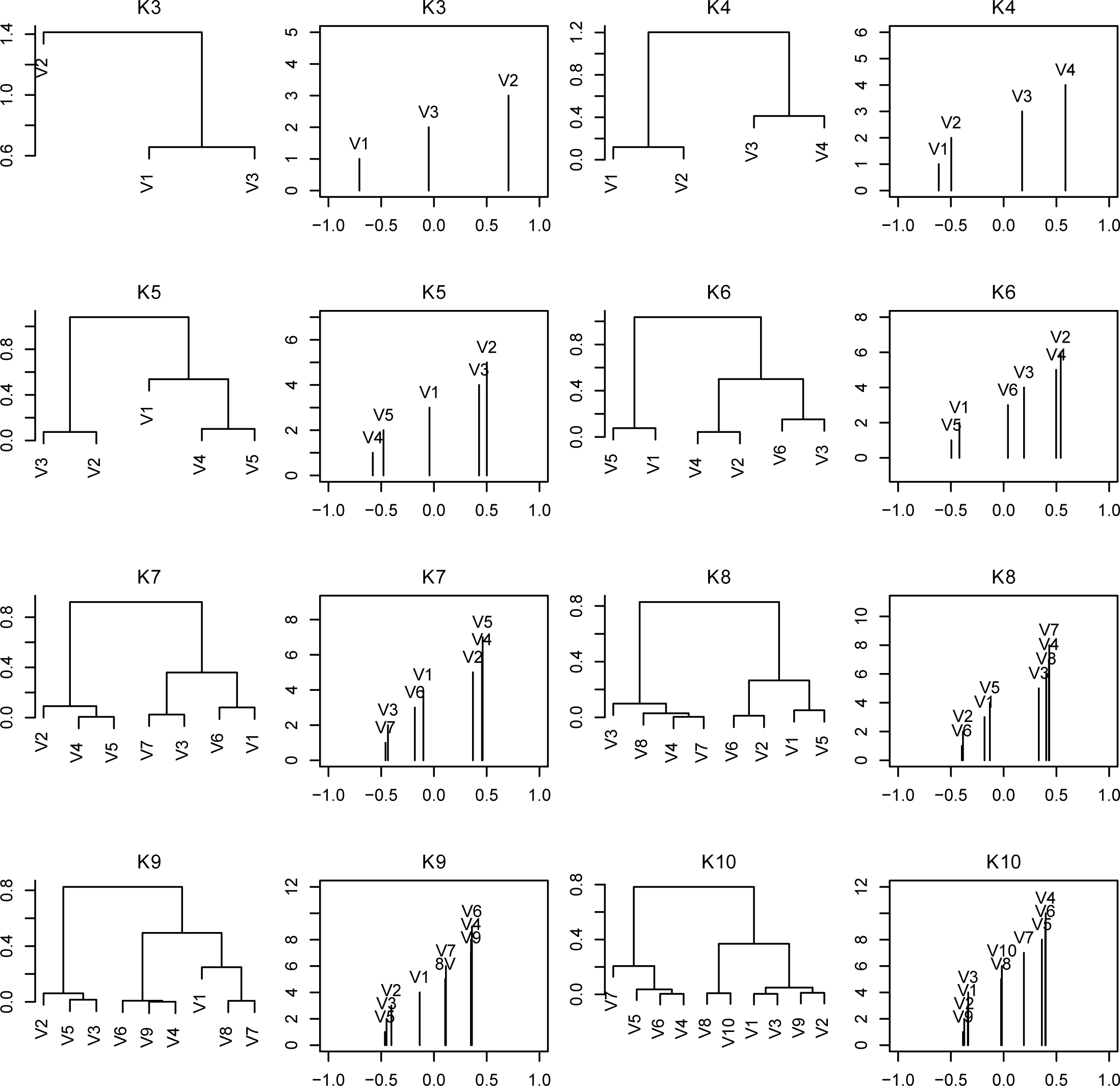
Hierarchical clustering and distance between classes on PC1. The relationship between clusters at each value of *k* is represented as both a dendrogram and as bar plot. The labels on the dendrogram (i.e., V1, V2, V3,…, V10) represent the original cluster assignment from *k*-means. The barplot to the right of each dendrogram depicts the elements of the eigenvector associated with the largest eigenvalue form PPKC. The labels above each line represent the original cluster assignment.

**Figure S5.**
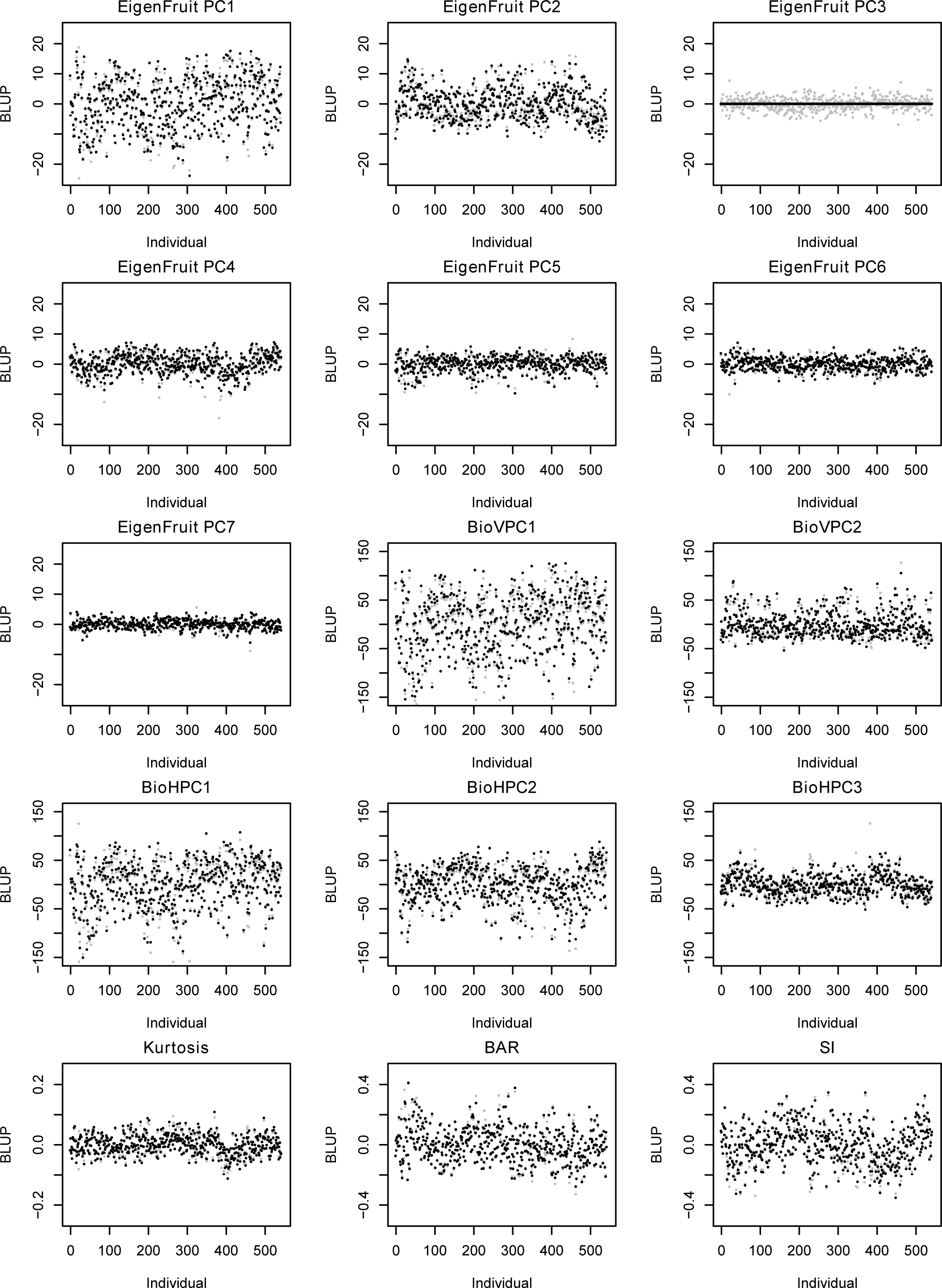
BLUPs for 15 selected features. For each plot, the X-axis is the index and the Y-axis is the BLUP value estimated from a linear mixed model. Grey points represent the mean feature value for each individual. Each point is the BLUP for a single genotype.

**Figure S6.**
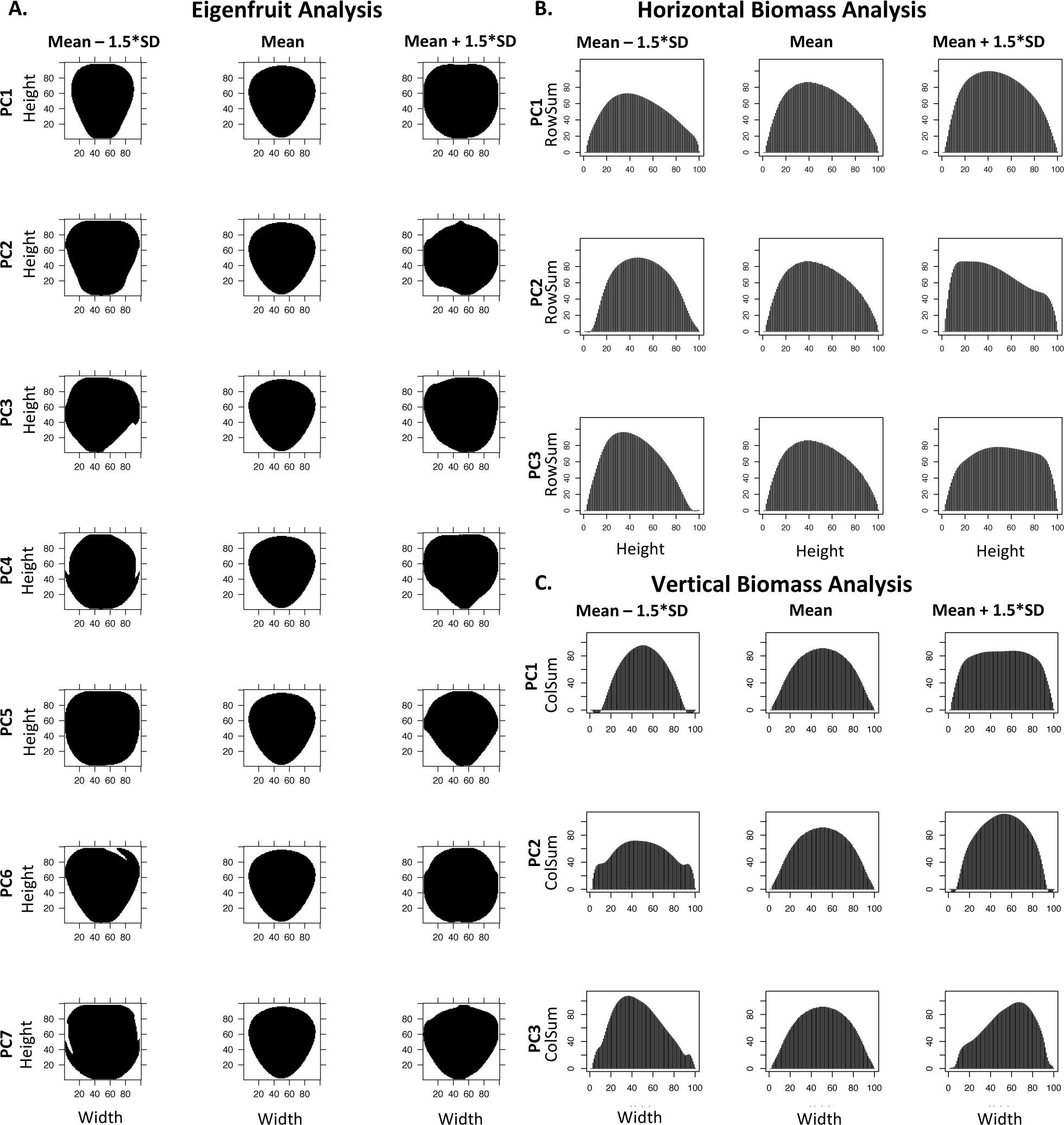
Effects of Eigenfruit, Vertical Biomass, and Horizontal Biomass Analyses. **(A)** Effects of PC [1, 7] from the Eigenfruit analysis on the mean shape (center column). Left column is the mean shape minus 1.5× the standard deviation. Right is the mean shape plus 1.5× the standard deviation. The horizontal axis is the horizontal pixel position. The vertical axis is the vertical pixel position. **(B)** Effects of PC [1, 3] from the Horizontal Biomass analysis on the mean shape (center column). Left column is the mean shape minus 1.5× the standard deviation. Right is the mean shape plus 1.5× the standard deviation. The horizontal axis is the vertical position from the image (height). The vertical axis is the number of activated pixels (RowSum) at the given veritcal position. **(C)** Effects of PC [1, 3] from the Vertical Biomass analysis on the mean shape (center column). Left column is the mean shape minus 1.5× the standard deviation. Right is the mean shape plus 1.5× the standard deviation. The horizontal axis is the horizontal position from the image (width). The vertical axis is the number of activated pixels (ColSum) at the given horizontal position.

**Figure S7.**
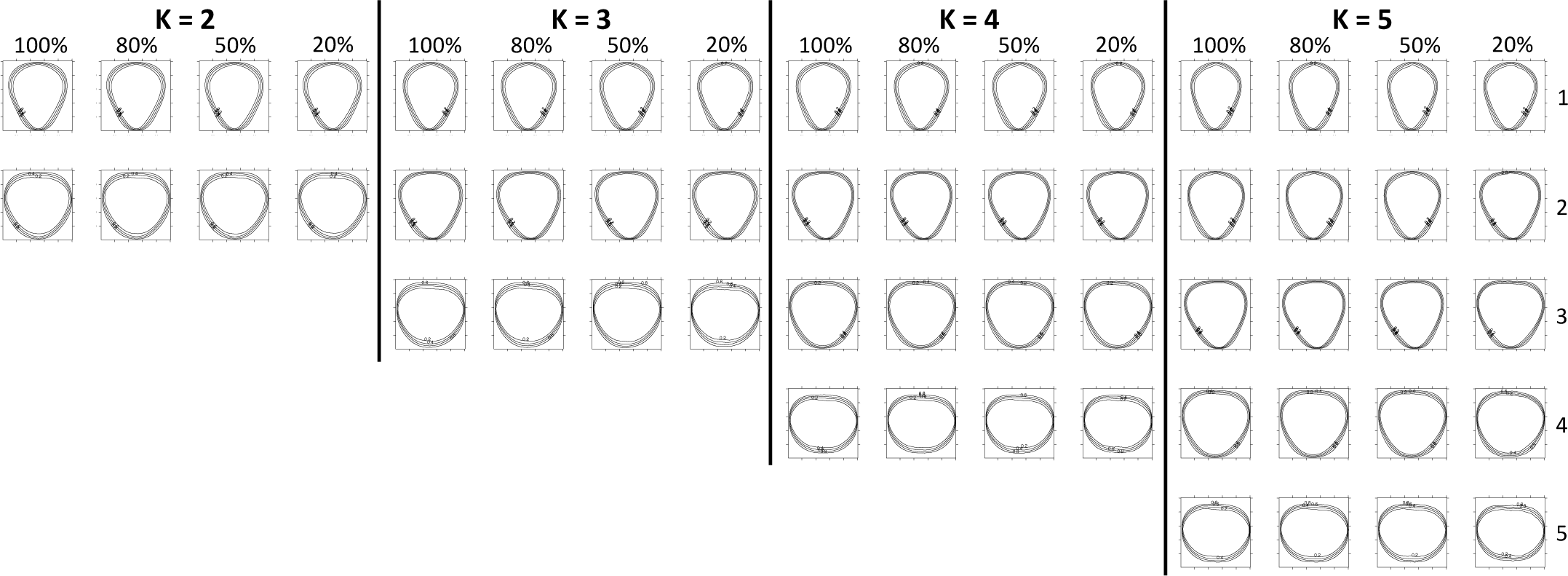
PPKC with variable sample size. Ordered centroids from *k* = 2 to *k* = 5 using different image sets for clustering. For all *k* = [2, 5], *k*-means clustering was performed using either 100, 80, 50%, or 20% of the total number of images; 6, 874, 5, 500, 3, 437, and 1, 374 respectively. Cluster position indicated on the right [1, 5].

## References

1. Duchesne A. Histoire naturelle des fraisiers. Didot le Jeune, Paris.; 1766.

2. Darrow GM. The strawberry. History, breeding and physiology. Holt, Rinehart and Winston, New York; 1966.

3. Edger PP, Poorten TJ, VanBuren R, Hardigan MA, Colle M, McKain MR, et al. Origin and evolution of the octoploid strawberry genome. Nature Genetics 2019 Mar;51(3):541–547.

4. Hardigan MA, Poorten TJ, Acharya CB, Cole GS, Hummer KE, Bassil N, et al. Domestication of Temperate and Coastal Hybrids with Distinct Ancestral Gene Selection in Octoploid Strawberry. The Plant Genome 2018;11(3):0.

5. Aharoni A. Gain and Loss of Fruit Flavor Compounds Produced by Wild and Cultivated Strawberry Species. The Plant Cell 2004 Nov;16(11):3110–3131.

6. Wang SY, Lewers KS. Antioxidant Capacity and Flavonoid Content in Wild Strawberries. Journal of the American Society for Horticultural Science 2007 Sep;132(5):629–637.

7. Diamanti J, Capocasa F, Balducci F, Battino M, Hancock J, Mezzetti B. Increasing Strawberry Fruit Sensorial and Nutritional Quality Using Wild and Cultivated Germplasm. PLoS ONE 2012 Oct;7(10):e46470.

8. Vallarino JG, de Abreu e Lima F, Soria C, Tong H, Pott DM, Willmitzer L, et al. Genetic diversity of strawberry germplasm using metabolomic biomarkers. Scientific Reports 2018 Dec;8(1).

9. Liao X, Li M, Liu B, Yan M, Yu X, Zi H, et al. Interlinked regulatory loops of ABA catabolism and biosynthesis coordinate fruit growth and ripening in woodland strawberry. Proceedings of the National Academy of Sciences 2018 Dec;115(49):E11542–E11550.

10. Whitaker VM, Hasing T, Chandler CK, Plotto A, Baldwin E. Historical Trends in Strawberry Fruit Quality Revealed by a Trial of University of Florida Cultivars and Advanced Selections. HortScience 2011 Apr;46(4):553–557.

11. Visa S, Cao C, Gardener BM, van der Knaap E. Modeling of tomato fruits into nine shape categories using elliptic fourier shape modeling and Bayesian classification of contour morphometric data. Euphytica 2014 Dec;200(3):429–439.

12. Migicovsky Z, Gardner KM, Money D, Sawler J, Bloom JS, Moffett P, et al. Genome to Phenome Mapping in Apple Using Historical Data. The Plant Genome 2016;9(2):0.

13. Turner SD, Ellison SL, Senalik DA, Simon PW, Spalding EP, Miller ND. An Automated Image Analysis Pipeline Enables Genetic Studies of Shoot and Root Morphology in Carrot (Daucus carota L.). Frontiers in Plant Science 2018 Nov;9.

14. Mathey MM, Mookerjee S, Gündüz K, Hancock JF, Iezzoni AF, Mahoney LL, et al. Large-Scale Standardized Phenotyping of Strawberry in RosBREED. Journal of the American Pomological Society 2013;p. 12.

15. Whitaker VM, Osorio LF, Hasing T, Gezan S. Estimation of Genetic Parameters for 12 Fruit and Vegetative Traits in the University of Florida Strawberry Breeding Population. Journal of the American Society for Horticultural Science 2012 Sep;137(5):316–324.

16. Antanaviciute L. Genetic mapping and phenotyping plant characteristics, fruit quality and disease resistance traits in octoploid strawberry (Fragaria× ananassa). PhD thesis, University of Reading; 2016.

17. Simpson MG. 9. In: Plant systematics Academic press; 2010. p. 494–508.

18. Ishikawa T, Hayashi A, Nagamatsu S, Kyutoku Y, Dan I, Wada T, et al. Classification of Strawberry Fruit Shape by Machine Learning. ISPRS - International Archives of the Photogrammetry, Remote Sensing and Spatial Information Sciences 2018 May;XLII-2:463–470.

19. dos Anjos RS, Marçal TdS, Carneiro P, Carneiro JEdS. New Proposals to Estimate Unbiased Selection Gain and Coefficient of Variation in Traits Evaluated Using Score Scales. Crop Science 2019;.

20. Mitry D, Zutis K, Dhillon B, Peto T, Hayat S, Khaw KT, et al. The accuracy and reliability of crowdsource annotations of digital retinal images. Translational Vision Science and Technology 2016;5(5):6–6.

21. Zhou N, Siegel ZD, Zarecor S, Lee N, Campbell DA, Andorf CM, et al. Crowdsourcing image analysis for plant phenomics to generate ground truth data for machine learning. PLoS Computational Biology 2018;14(7):e1006337.

22. Chollet F, Allaire JJ. Deep Learning with R. 1st ed. Greenwich, CT, USA: Manning Publications Co.; 2018.

23. Achcar F, Camadro JM, Mestivier D. AutoClass@ IJM: a powerful tool for Bayesian classification of heterogeneous data in biology. Nucleic acids research 2009;37(suppl_2):W63–W67.

24. Cheverud JM, Buikstra JE. Quantitative genetics of skeletal nonmetric traits in the rhesus macaques on Cayo Santiago. II. Phenotypic, genetic, and environmental correlations between traits. American Journal of Physical Anthropology 1981;54(1):51–58.

25. Agresti A. Analysis of ordinal categorical data, vol. 656. John Wiley & Sons; 2010.

26. Montesinos-López OA, Montesinos-López A, Pérez-Rodríguez P, de los Campos G, Eskridge K, Crossa J. Threshold Models for Genome-Enabled Prediction of Ordinal Categorical Traits in Plant Breeding. G3: Genes, Genomes, Genetics 2015 Feb;5(2):291–300.

27. Montesinos-López OA, Montesinos-López A, Crossa J, Burgueño J, Eskridge K. Genomic-Enabled Prediction of Ordinal Data with Bayesian Logistic Ordinal Regression. G3: Genes, Genomes, Genetics 2015 Oct;5(10):2113–2126.

28. Fresnedo-Ramírez J, Famula TR, Gradziel TM. Application of a Bayesian ordinal animal model for the estimation of breeding values for the resistance to Monilinia fruticola (G. Winter) Honey in progenies of peach [Prunus persica (L.) Batsch]. Breeding Science 2017;p. 16027.

29. Hearn DJ. Shape analysis for the automated identification of plants from images of leaves. Taxon 2009 Aug;58(3):934–954.

30. Fu G, Berg A, Das K, Li J, Li R, Wu R. A statistical model for map-ping morphological shape. Theoretical Biology and Medical Modelling 2010;7(1):28.

31. Balduzzi M, Binder BM, Bucksch A, Chang C, Hong L, Iyer-Pascuzzi AS, et al. Reshaping Plant Biology: Qualitative and Quantitative Descriptors for Plant Morphology. Frontiers in Plant Science 2017 Feb;08.

32. Tanksley SD. The genetic, developmental, and molecular bases of fruit size and shape variation in tomato. The plant cell 2004;16(suppl 1):S181–S189.

33. Monforte AJ, Diaz A, Caño-Delgado A, van der Knaap E. The genetic basis of fruit morphology in horticultural crops: lessons from tomato and melon. Journal of Experimental Botany 2013 Aug;65(16):4625–4637.

34. Lynch M, Walsh B, et al. Genetics and analysis of quantitative traits, vol. 1. Sinauer Sunderland, MA; 1998.

35. Goddard M, Hayes B. Genomic selection. Journal of Animal breeding and Genetics 2007;124(6):323–330.

36. Heffner EL, Sorrells ME, Jannink JL. Genomic selection for crop improvement. Crop Science 2009;49(1):1–12.

37. Resende MF, Muñoz P, Resende MD, Garrick DJ, Fernando RL, Davis JM, et al. Accuracy of genomic selection methods in a standard data set of loblolly pine (Pinus taeda L.). Genetics 2012;190(4):1503–1510.

38. Xiao H, Jiang N, Schaffner E, Stockinger EJ, Van Der Knaap E. A retrotransposon-mediated gene duplication underlies morphological variation of tomato fruit. Science 2008;319(5869):1527–1530.

39. Wu S, Zhang B, Keyhaninejad N, Rodríguez GR, Kim HJ, Chakrabarti M, et al. A common genetic mechanism underlies morphological diversity in fruits and other plant organs. Nature Communications 2018 Dec;9(1).

40. Han K, Jeong HJ, Yang HB, Kang SM, Kwon JK, Kim S, et al. An ultra-high-density bin map facilitates high-throughput QTL mapping of horticultural traits in pepper (Capsicum annuum). DNA Research 2016;23(2):81–91.

41. Chunthawodtiporn J, Hill T, Stoffel K, Van Deynze A. Quantitative trait loci controlling fruit size and other horticultural traits in bell pepper (Capsicum annuum). The Plant Genome 2018;11(1).

42. White AG, Alspach PA, Weskett RH, Brewer LR. Heritability of fruit shape in pears. Euphytica 2000 Mar;112(1):1–7.

43. Prashar A, Hornyik C, Young V, McLean K, Sharma SK, Dale MFB, et al. Construction of a dense SNP map of a highly heterozygous diploid potato population and QTL analysis of tuber shape and eye depth. Theoretical and Applied Genetics 2014 Oct;127(10):2159–2171.

44. Lerceteau-Köhler E, Moing A, Guérin G, Renaud C, Petit A, Rothan C, et al. Genetic dissection of fruit quality traits in the octoploid cultivated strawberry highlights the role of homoeo-QTL in their control. Theoretical and Applied Genetics 2012;124(6):1059–1077.

45. Tanabata T, Shibaya T, Hori K, Ebana K, Yano M. SmartGrain: highthroughput phenotyping software for measuring seed shape through image analysis. Plant physiology 2012;160(4):1871–1880.

46. Kuhl FP, Giardina CR. Elliptic Fourier features of a closed contour. Computer Graphics and Image Processing 1982;18(3):236–258.

47. Chitwood DH, Ranjan A, Martinez CC, Headland LR, Thiem T, Kumar R, et al. A Modern Ampelography: A Genetic Basis for Leaf Shape and Venation Patterning in Grape. Plant Physiology 2014 Jan;164(1):259–272.

48. Li M, An H, Angelovici R, Bagaza C, Batushansky A, Clark L, et al. Topological Data Analysis as a Morphometric Method: Using Persistent Homology to Demarcate a Leaf Morphospace. Frontiers in Plant Science 2018 Apr;9.

49. Chitwood DH, Otoni WC. Morphometric analysis of Passiflora leaves: the relationship between landmarks of the vasculature and elliptical Fourier descriptors of the blade. GigaScience 2017 Jan;6(1).

50. Li M, Frank MH, Coneva V, Mio W, Chitwood DH, Topp CN. The persistent homology mathematical framework provides enhanced genotype-to-phenotype associations for plant morphology. Plant physiology 2018;177(4):1382–1395.

51. Gower JC. Generalized procrustes analysis. Psychometrika 1975;40(1):33–51.

52. Bookstein FL. Landmark methods for forms without landmarks: morphometrics of group differences in outline shape. Medical Image Analysis 1997 Apr;1(3):225–243.

53. Klingenberg CP, Leamy LJ. Quantitative Genetics of Geometric Shape in the Mouse Mandible. Evolution 2001;55(11):2342–2352.

54. Langlade NB, Feng X, Dransfield T, Copsey L, Hanna AI, Thébaud C, et al. Evolution through genetically controlled allometry space. Proceedings of the National Academy of Sciences 2005;102(29):10221–10226.

55. Bensmihen S, Hanna AI, Langlade NB, Micol JL, Bangham A, Coen ES. Mutational spaces for leaf shape and size. Hfsp Journal 2008;2(2):110–120.

56. Manacorda CA, Asurmendi S. Arabidopsis phenotyping through geometric morphometrics. GigaScience 2018;7(7):giy073.

57. Sirovich L, Kirby M. Low-dimensional procedure for the characterization of human faces. Journal of the Optical Society of America 1987 Mar;4(3):519.

58. Turk MA, Pentland AP. Face recognition using eigenfaces. In: Proceedings. 1991 IEEE Computer Society Conference on Computer Vision and Pattern Recognition; 1991. p. 586–591.

59. Horgan GW, Talbot M, Davey JC. Use of statistical image analysis to discriminate carrot cultivars. Computers and Electronics in Agriculture 2001;31(2):191–199.

60. Horgan GW. The statistical analysis of plant part appearance—a review. Computers and Electronics in Agriculture 2001;31(2):169–190.

61. Ehsanirad A. Plant classification based on leaf recognition. International Journal of Computer Science and Information Security 2010;8(4):78–81.

62. Rodrigo R, Samarawickrame K, Mindya S. An Intelligent Flower Analyzing System for Medicinal Plants. Conference on Computer Graphics, Visualization and Computer Vision 2013;p. 4.

63. Feldmann MJ. Classification and Quantification of Strawberry Fruit Shape Data; 2019, http://dx.doi.org/10.5281/zenodo.3365715.

64. Feldmann MJ. 2DShapeDescription; 2019, https://github.com/mjfeldmann/2DShapeDescription.

65. Lloyd SP. Least squares quantization in pcm. IEEE Transactions on Information Theory 1982;28:129–137.

66. Evanno G, Regnaut S, Goudet J. Detecting the number of clusters of individuals using the software STRUCTURE: a simulation study. Molecular Ecology 2005;14(8):2611–2620.

67. Christensen RHB. ordinal—Regression Models for Ordinal Data; 2019, r package version 2019. 3–9. http://www.cran.r-project.org/package=ordinal/.

68. Genuer R, Poggi JM, Tuleau-Malot C. VSURF: an R package for variable selection using random forests. The R Journal 2015;7(2):19–33.

69. Bates D, Mächler M, Bolker B, Walker S. Fitting Linear Mixed-Effects Models Using lme4. Journal of Statistical Software 2015;67(1):1–48.

70. Bernardo R, Thompson AM. Germplasm architecture revealed through chromosomal effects for quantitative traits in maize. The plant Genome 2016;9(2).

71. Voth V, Bringhurst RS. Strawberry plant called Chandler; 1984, uS Patent App. 06/452,699.

72. Voth V, Shaw DV, Bringhurst RS. Strawberry plant called Camarosa; 1994, uS Patent App. 08/041,742.

73. Suenaga T, Imamura Y, Maeda K, Yamada T, Takamatsu M. The workloads of farmers who sort and pack strawberries in accordance with standards of shipment and their awareness of standards of shipment. Journal of the Japanese Association of Rural Medicine 1989;38(4):895–907.

74. Sonnenschein A, VanderZee D, Pitchers WR, Chari S, Dworkin I. An image database of Drosophila melanogaster wings for phenomic and biometric analysis. GigaScience 2015;4(1):25.

75. Pincot DD, Poorten TJ, Hardigan MA, Harshman JM, Acharya CB, Cole GS, et al. Genome-wide association mapping uncovers Fw1, a dominant gene conferring resistance to Fusarium wilt in strawberry. G3: Genes, Genomes, Genetics 2018;8(5):1817–1828.

76. Jiang N, Gao D, Xiao H, Van Der Knaap E. Genome organization of the tomato sun locus and characterization of the unusual retrotransposon Rider. The Plant Journal 2009;60(1):181–193.

77. Frary A, Nesbitt TC, Frary A, Grandillo S, Van Der Knaap E, Cong B, et al. fw2. 2: a quantitative trait locus key to the evolution of tomato fruit size. Science 2000;289(5476):85–88.

78. Liu J, Van Eck J, Cong B, Tanksley SD. A new class of regulatory genes underlying the cause of pear-shaped tomato fruit. Proceedings of the National Academy of Sciences 2002;99(20):13302–13306.

79. Rodríguez GR, Muños S, Anderson C, Sim SC, Michel A, Causse M, et al. Distribution of SUN, OVATE, LC, and FAS in the tomato germplasm and the relationship to fruit shape diversity. Plant physiology 2011;156(1):275–285.

80. Rodríguez GR, Kim HJ, Van Der Knaap E. Mapping of two suppressors of OVATE (sov) loci in tomato. Heredity 2013;111(3):256.

81. Lande R, Thompson R. Efficiency of marker-assisted selection in the improvement of quantitative traits. Genetics 1990;124(3):743–756.

82. Wang F. SIOX plugin in ImageJ: area measurement made easy. UV4 Plants Bulletin 2017 Feb;2:37–44.

83. Schneider CA, Rasband WS, Eliceiri KW. NIH Image to ImageJ: 25 years of image analysis. Nature Methods 2012 Jul;9(7):671–675.

84. Schindelin J, Arganda-Carreras I, Frise E, Kaynig V, Longair M, Pietzsch T, et al. Fiji: an open-source platform for biological-image analysis. Nature methods 2012;9(7):676.

85. R Core Team. R: A Language and Environment for Statistical Computing. R Foundation for Statistical Computing, Vienna, Austria; 2019, https://www.R-project.org/.

86. Urbanek S. jpeg: Read and write JPEG images; 2014, r package version 0.1-8.

87. Ooms J. magick: Advanced Graphics and Image-Processing in R; 2018, r package version 2.0.

88. Bonhomme V, Picq S, Gaucherel C, Claude J. Momocs: Outline Analysis Using R. Journal of Statistical Software 2014;56(13):1–24.

89. Rosseel Y. lavaan: An R Package for Structural Equation Modeling. Journal of Statistical Software 2012;48(2):1–36.

90. Schreiber JB, Nora A, Stage FK, Barlow EA, King J. Reporting Structural Equation Modeling and Confirmatory Factor Analysis Results: A Review. The Journal of Educational Research 2006 Jul;99(6):323–338.

91. Venables WN, Ripley BD. Modern Applied Statistics with S. Fourth ed. New York: Springer; 2002. ISBN 0-387-95457-0.

92. Meyer D, Dimitriadou E, Hornik K, Weingessel A, Leisch F. e1071: Misc Functions of the Department of Statistics, Probability Theory Group (Formerly: E1071), TU Wien; 2019, r package version 1.7-0.1.

